# Removal of behavioural and electrophysiological signs of chronic pain by *in vivo* microsections of rat somatosensory cortex with parallel X-ray microbeams

**DOI:** 10.1101/528539

**Authors:** Antonio G. Zippo, Gloria Bertoli, Maria Pia Riccardi, Maurizio Valente, Elke Bräuer-Krisch, Gian Carlo Caramenti, Herwig Requardt, Veronica del Grosso, Paola Coan, Alberto Bravin, Gabriele E. M. Biella

**Affiliations:** Institute of Molecular Bioimaging and Physiology, Consiglio Nazionale delle Ricerche, Segrate (Milan), Italy.; Earth and Environment Department and Arvedi Laboratory, Università degli Studi di Pavia, Pavia, Italy; ESRF, ID17, Rue de Martyrs, Grenoble, France; Ludwig-Maximilians-Universität, Department of Clinical Radiology and Department of Physics, München, Deutschland

## Abstract

Chronic pain (CP) is a condition characterized by a wide spectrum of clinical signs and symptoms, missing a sound modelling at the neuronal network scale. Recently, we presented a general theory showing common electrophysiological traits in different CP rat models, i.e. a collapse of relevant functional connectivity network properties, such as modularity, in the somatosensory thalamo-cortical (TC) network. In this work, we preliminary investigated by an *in silico* accurate simulator of the six-layer mammalian cortical networks that evidenced the crucial collapse of network modularity in CP simulated conditions and the consequent reduction of network adaptive processes. On this track, in studies on CP experimental animals affected by sciatic nerve multiple ligature (Bennett-Xie model), by synchrotron-generated X-ray microbeam (MB) irradiations (7 parallel beams, 100um width), we targeted *in vivo* the CP involved hindlimb somatosensory projection cortex that, because of the doses radiation (360 Gy, peak at each beam), non-invasively produced fast and precise tissue destruction along the 7 beam projections. These parcellated the cortical tissue and restored the cortical network statistics related to modularity and information processing efficiency as evidenced from post irradiation *in vivo* electrophysiological recordings. In addition, by MB treatment there was an ensuing removal of behavioral signs of allodynia and hyperalgesia accompanied by recovered normal gait schemes yet preserving the normal sensory thresholds of the experimental rats up to three months after the MB irradiation. Finally, novel and unprecedented therapeutic appraisals for CP are devised.

**Significance Statement:** Chronic pain (CP) is an excruciating condition with severe effects on patients’ life. Apart from many clinical and experimental studies no current theory on CP is generally accepted. Recently, we proposed a general theory of CP in experimental animals as characterized by strong alteration of the connections among neurons in different brain regions. We show here on *in silico* simulations that specific connectivity changes in the somatosensory cortex recover the lost functional integrity. Concurrently, in experimental animals, we re-modulated, *in vivo*, some anatomical connections of the somatosensory cortex by extremely thin synchrotron generated X-ray microbeam irradiations. The resulting behavioral and electrophysiological signs of CP disappeared yet maintaining normal sensory responses. No adverse or pathological effects on blank animals were observable.

## Introduction

Chronic Pain (CP) is a multiform sensory disorder (Basbaum et al. 2009; Descalzi et al. 2015) combining plural signs and symptoms with genetic and epigenetic (Liang et al. 2015) traits. Despite the stream of works dedicated to many aspects of CP (Apkarian, Baliki, and Geha 2009; Melzack 2001), a general model of CP is not yet available at the scales of neuronal networks. In a recent appeared novel interpretation, different experimental models of CP shared a common set of anomalies of the neuronal network connectivity in the thalamocortical axis (A.G. Zippo et al. 2016). In a recent model, CP has been conformed to an altered connectivity disorder where the somatosensory thalamocortical network exhibits reduced capabilities of information processing along with weakened information transfer and overall topological network degradation. As a whole, these conditions concur to generate stereotyped network configurations, which seemingly appear as anatomo-functional counterparts of the continuous percept of pain, a picture recoverable from all the examined CP models. The model is theoretically sustained by the fact that persistent acute pain elicits massive hyperactivation of thalamocortical neurons strongly modifying the synaptic loads (Vierck et al. 2013; Mansour et al. 2014), which in turn dramatically alter the functional organization of the involved neuronal networks (Spisák et al. 2017; Gustin et al. 2012). In this dramatic context, actual conventional CP therapies do not appear adapt to relocate or repair within original limits the disordered network connectivity. This may be potentially due to many factors such as the relocation of countless functional multiscale parameters (e.g. ion channel typing, synaptic dynamics or again plastic responses to environmental requirements). Novel putative strategies appear, thus, long awaited eventually changing the current therapeutic approaches, those being pharmacological or neurosurgical (Price et al. 2018). With the assumption that CP appears too much a complex multiscale disorders, we conjectured that a widely encompassing intervention focussed on the physical re-arrangement of the involved thalamocortical network architectures could prompt concurrent modifications in the information processing leading to sensory re-conversion back to match control conditions (Francis and Song 2011).

From an anatomo-functional point of view, the dense cortical somatotopic projections are characterized by dynamic borders maintained on a delicate balance of excitation/inhibition through the cortical circuits, especially with horizontal connections (B A Johnson and Frostig 2016; Brett A Johnson and Frostig 2018; Négyessy et al. 2013). These refined networks, designing a functional modularity in the cortical somatotopic projection fields, undergo a failure in CP with integer contiguity loss and a derangement of projection field independence. This loss of modularity (the edge fraction belonging to the given groups minus the expected fraction in comparison to randomly distributed edges) may have strong implications on the circuitry dynamics (Darian-Smith and Gilbert 1994). A strategy based on an “artificial” restoration of module-like architecture could retrace a design more apt to better responding to the functional needs of the local cortical circuits. The hypothesis is based on a previous paper by our group, proposing a novel theoretic context for chronic pain. Namely, that CP, analyzed by simultaneous thalamic and cortical massive microelectrode neuronal recordings, could be described as a loss of functional connectivity and reduced thalamo-cortical information transfer. Here, by a large-scale cortical six-layer column simulator, we investigated the pivotal role of intercolumnar connectivity. Operatively, such connections are hard to modulate though. However, by Synchrotron driven high-dose (360 Greys, Gy) X-ray microbeams (MB) irradiation into the brain *in vivo* (i.e. X-rays generated by a synchrotron light source are collimated into wafers of parallel microbeams few tens of microns wide, separated by a few hundred microns) we induced non-invasive thin cuts (e.g. 100 µm wide) within the primary somatosensory cortex in order to induce connection discontinuities, orthogonal to the cortical surface (Babadi et al. 2014). Concurrent researches on other pathologies, such as epilepsy, have shown the successful technical application and feasibility of this procedure on living rats by the delivery of microbeams in hippocampal regions (Pouyatos et al. 2016; Romanelli et al. 2013). Due to the technical features of the X ray emitting source and the anatomical constraints mainly horizontal connections have thus been addressed, though their still unclear role in CP, just to start an experimental series aiming to explore how tiniest X-ray guided brain network microlesions in “closed sky” conditions (with no actual classical surgical invasive skull opening technique) could achieve functional crucial results able to manipulate sensory functional connectivity and the related neural processing.

In experimental animal sessions, these behavioral and electrophysiological results suggest novel therapeutic approaches potentially reforming current outcomes.

## Materials and Methods

### Ethical Statement

All the animals (28 albino Sprague-Dawley male rats 225-250g) have been used in accordance to the Italian and European Laws on animal treatment in Scientific Research (Italian Bioethical Committee, Law Decree on the Treatment of Animals in Research, 27 Jan 1992, No. 116, guideline n. 86/609/ European Community). The research was approved by the Ministry of Health and classified as “Biella 3/2011 and further as “Biella 1/2014” into the files of the Ethical Committee of the University of Milan, Italy and 14_ethax16 for the experiments at the ESRF ID17 beamline (European Synchrotron Radiation Facility, ESRF, Grenoble, France).

## Results

The results report a multianalytic picture of the effects of high-dose sevenfold X-ray microbeams (MBs) irradiation through the intact skull on the somatosensory cortex of experimental rats *in vivo* anesthetized with a model of neuropathic Chronic Pain (CP, Figure 1A-F) and control (CR).

**Figure 1.**
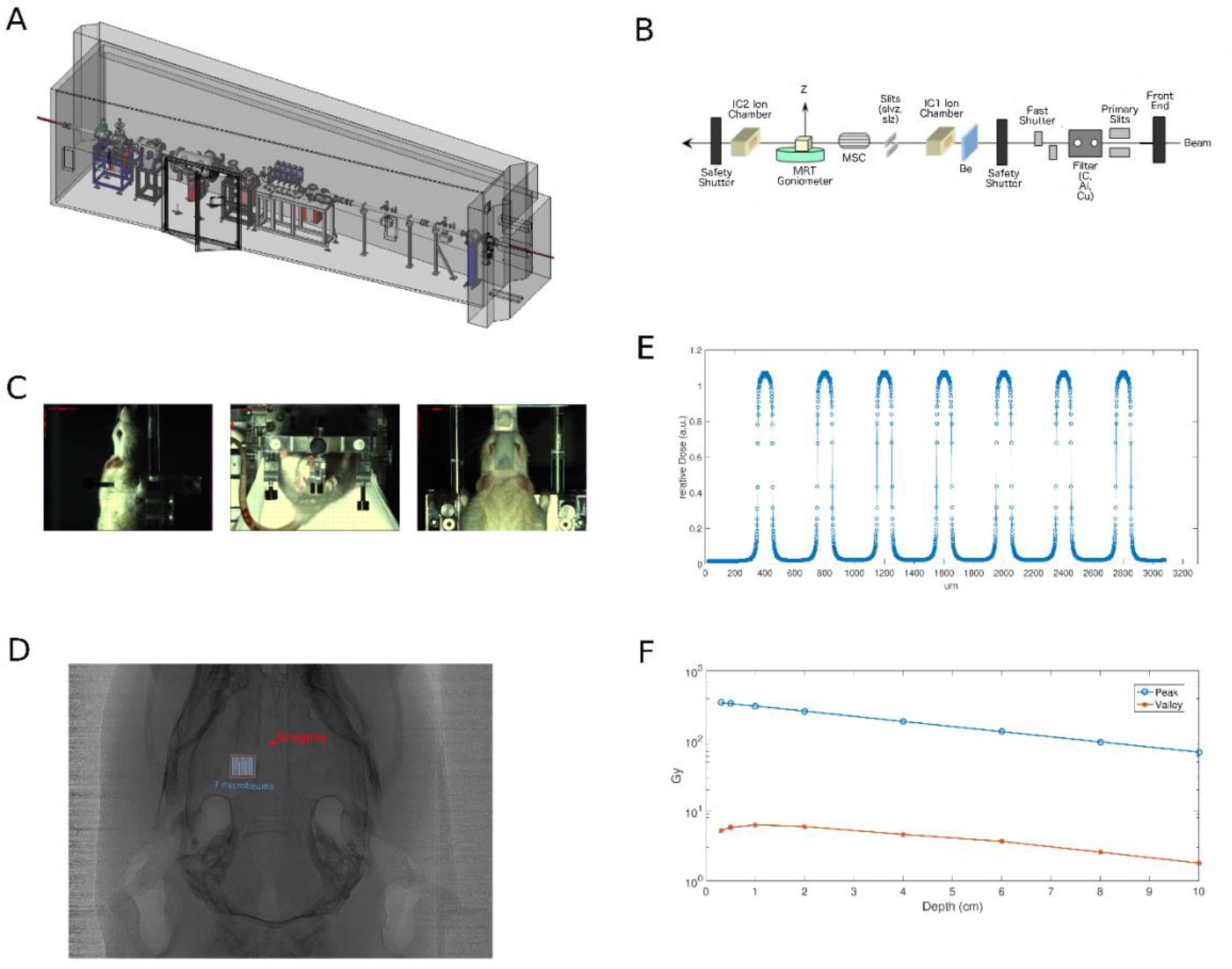
Experimental setup of the microbeams irradiation on rat brain. (A) Schematic of the ESRF ID17 synchrotron line; the elements are not to scale. (B) Schematic of beam optoprocessing along the line. The beam issued by the wiggler is first shaped by the slits, filtered by a series of C, Al, Cu filters which removed the low energy components of the spectrum, monitored by an ionization chamber and spatially fractionated by the multi slit collimator (MSC); the sample is placed on the goniometer (C) Stereotactic placement of the rat. (D) X-ray imaging of the rat for stereotactic targeting, the Bregma point is highlighted by the red spot and the targeted S1 area, by the seven (blue) beams, is labeled by the red square. (E) Spatial distribution of the seven microbeams with peak and valley relative doses. (F) Dose deposited by the X-ray microbeams vs the tissue depth.

### Early computational inspections

Preliminary, *in silico* exploratory analyses offered the possibility to evaluate the effects of guided connectivity interventions on functional and topological substrates of the cortical network. Specifically, a comparable network of the hindlimb six-layer primary somatosensory cortex (2.4mm × 3mm, ∼1.5 millions of neurons) was simulated by means of the “Vertex Simulator”, a Matlab (The MathWorks, Inc., Natick, Massachusetts, United States) toolbox for large-scale cortical models (Tomsett et al. 2015). Our working hypothesis was that, since network adaptivity is directly correlated with network modularity, by artificially increasing the number of network modules, the network capability to adapt to the chronic nociceptive conditions should increase accordingly. Novel network modules were generated by reducing horizontal connections resulting in the best strategy to jointly maximize the number of novel modules and minimizing the number of removed connections. To this purpose, we reproduced virtual recording sessions, a facility provided by the toolbox, of a small subset of neurons (about one hundred, comparable to a conventional multielectrode experimental recording session) in four different conditions: i) a physiological state where activity is ignited by stochastic inputs that replicated relevant cortical dynamics; ii) a nociceptive state simulated by five to tenfold stronger inputs; iii) a physiological state (as in i)) characterized by weakened horizontal connection (*horizontal transection*, HT); iv) a nociceptive state (as in iii)) characterized by weakened horizontal connection.

Primarily, the influence of the relative amount of removed horizontal connections was evaluated on the complex network statistics of interest that were the Clustering Coefficient (C), the Efficiency (E) and the Modularity (Q), direct indicators of the network integrity. From 0 (original network) to 100 (all horizontal connection removed) with a resolution step of 1, we repeated the simulations 100 times to face the inherent randomness of the model. We found that all network statistics peaked between 7 and 9% (Figure 2A-C). Thus, we chose 8% as the relative quantity of randomly remove horizontal connections from the original network model in order to maximize modularity, efficiency and clustering coefficient.

**Figure 2.**
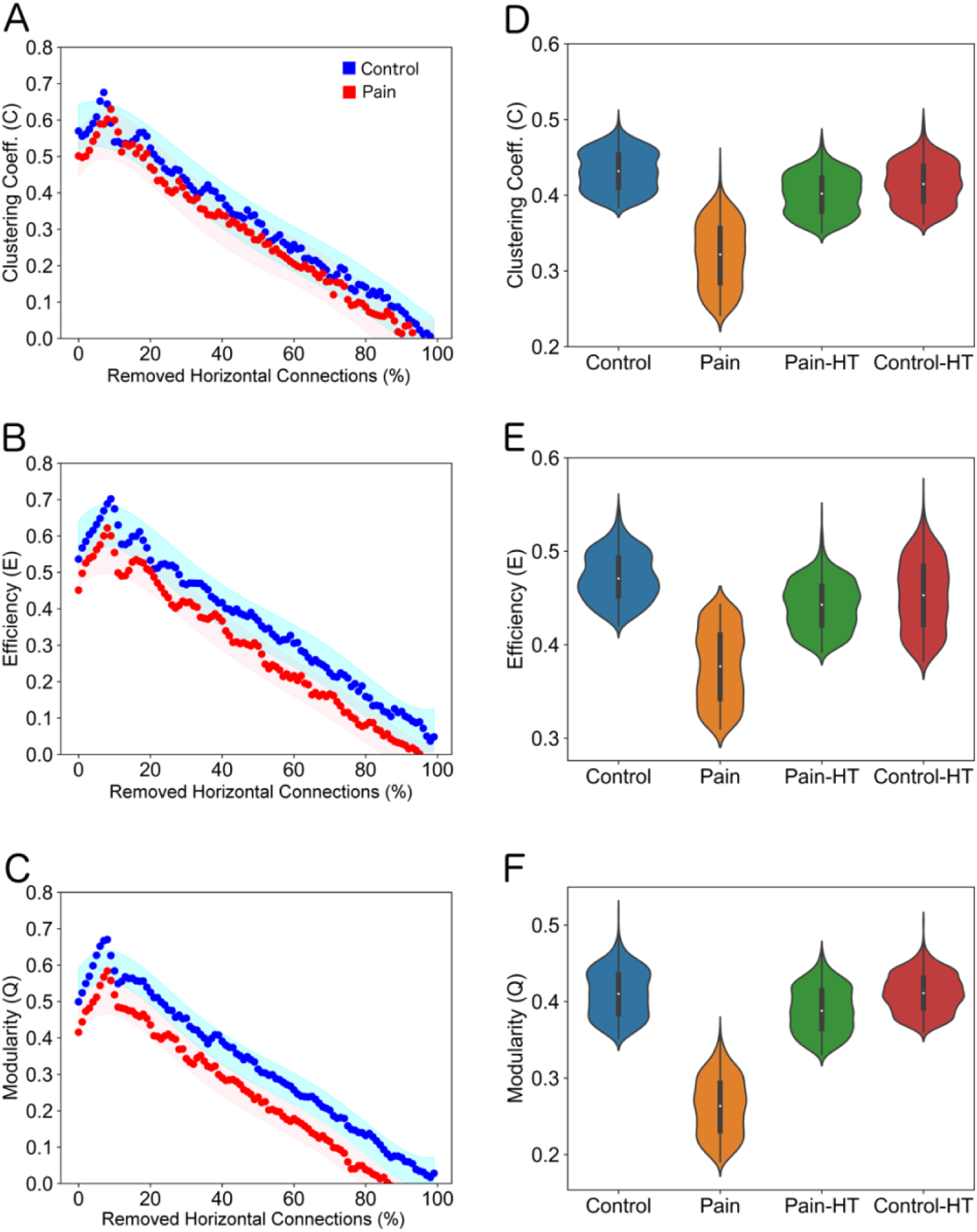
Results of cortical network simulations. The effects of the horizontal connection removal on the clustering coefficient (A), the efficiency (B) and the modularity (C). All plots clearly show peaks of values around 7-9%. By removing the 8% of the horizontal connections we found a complete restoring of the three statistics above strongly affected by noxious conditions (a simulation of sustained pain state). Transection of horizontal connections keep unaffected network statistics of the control dynamics. In (A)-(C), each point indicates the output of a simulation (N = 200 for each plot). Red points indicate the pain condition while blue point refer to control. Related bands represent the 95% confidence intervals. In (D)-(F), violin plots represent the distribution of each set of simulations.

Results of such simulations showed that network modularity collapsed in nociceptive conditions (P < 0.001, N = 1000, Wilcoxon non-parametric ranksum test with Bonferroni correction) in accordance with experimental works(A.G. Zippo et al. 2016). Modularity in networks with weakened horizontal connections was increased in comparison to nociceptive states (see Figure 2F; “Pain-HT”: P < 0.001, N = 1000; “Control-HT”: P < 0.001, N = 1000, ranksum tests) while unaltered when applied in the normal state (P = 0.362, N = 1000, ranksum test). Likewise, an incremental pattern was found for relevant network statistics like the clustering coefficient (Figure 2D; “Pain-HT”: P < 0.001, N = 1000; “Control-HT”: P < 0.001, N = 1000, ranksum tests) and the network efficiency (Figure 2E; “Pain-HT”: P < 0.001, N = 1000; “Control-HT”: P < 0.001, N = 1000, ranksum tests) suggesting that, in the simulated conditions, a complete restoring of the normal topological properties was achievable by opportune modulation of the horizontal connectivity.

Therefore, in silico results supported a non-invasive intervention in which the ablation of a limited percentage of the cortical horizontal connections could provide a topological recovering of non-noxious states.

### Sensory tests: thresholds and latencies

According to literature (Millan et al. 1987), young rats underwent a transitory decline of the physiological body weight increment after the chronic sciatic ligature, a trend also confirmed in animals treated with X-ray microbeams (MB, Figure 3A).

**Figure 3.**
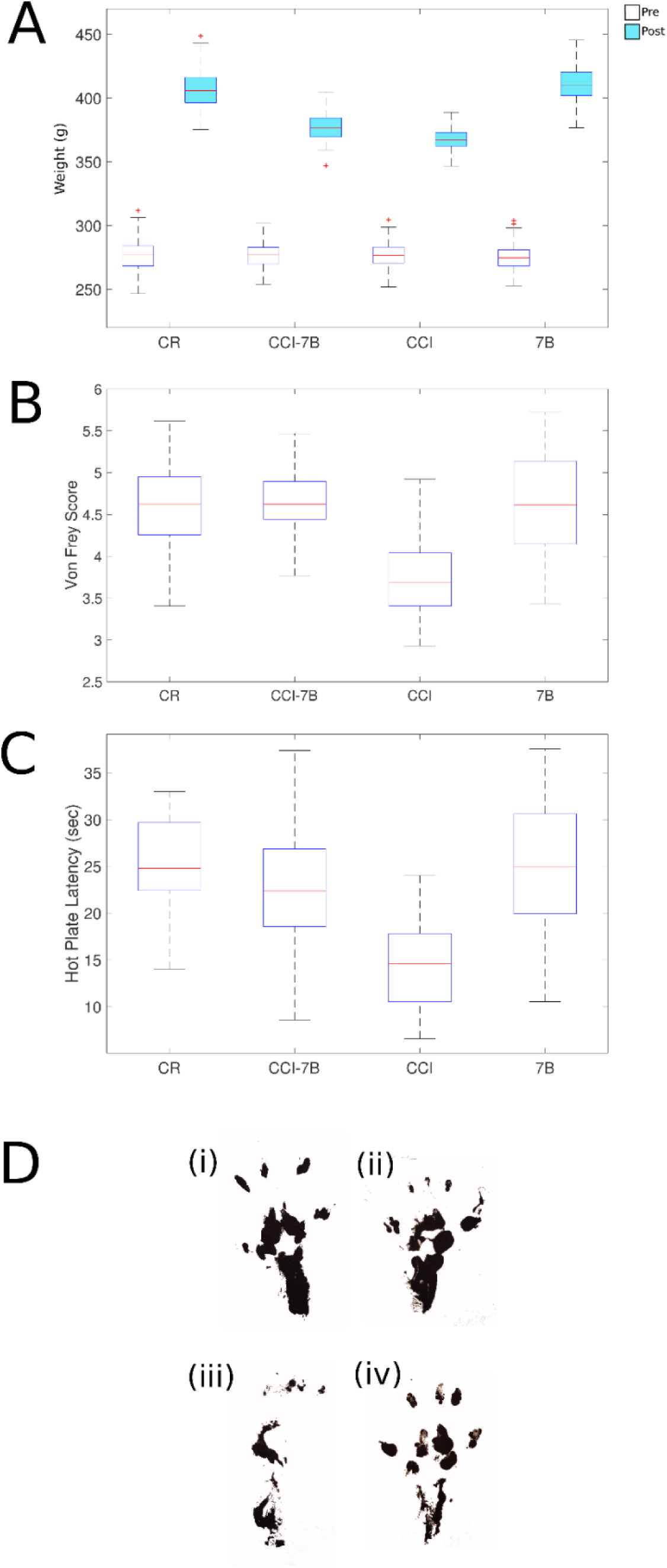
Behavioral signs of remission. (A) Box-Whisker plots of the body weight distribution in the four different groups during the pre-treatment (white) and post-treatment (cyan) stages. Box-Whisker plots of the von Frey (B) and the Hot plate test (C) scores. (D) Footprints of CR (i), CCI-MB (ii), CCI (iii) and CR-MB (iv).

In order to assess the effective emergence of hyperalgesia and allodynia in the CP experimental animals we performed the von Frey and the Hot Plate tests (Figure 3B-C). Indeed, the Von Frey score (Figure 3B), that presents important central responses (Ziegler et al. 1999), was reduced from an average of 4.73 ± 0.53 (standard deviation) in Control (CR) to 3.81 ± 0.41 in Chronic Constriction Injury (CCI) rats (P < 0.001, N = 28, non-parametric Wilcoxon ranksum test). The CCI-MB-treated rats showed instead a recovery from both allodynic and hyperalgesic conditions with an increase toward control values of 4.68 ± 0.55 after the MB treatment (CCI-MB) (P = 0.004, N = 28, ranksum test), statistically indiscernible from MB-treated control (CR-MB) (P = 0. 643, N = 28, ranksum test). Qualitatively, the videos S1-4 show the postural changes occurred in an exemplar CR animal before (Movie 1) and after (Movie 4) the MB treatment and in exemplar CCI animal before (Movie 2) and after (Movie 3) the MB treatment.

Results of the Hot Plate test, equally presenting important central effects(Gårdmark, Höglund, and Hammarlund-Udenaes 1998), showed (Figure 3C) a reduced latency in the CCI group (P < 0.001, N = 28, ranksum test) with retracted paws, reduced movement spans, and asymmetric posture in resting and walking schemes, restored in the MB treated animals (P = 0. 401, N = 28, ranksum test). Also, in this test the MB intervention did not significantly affect control (CR-MB) rats (P = 0. 873, N = 28, ranksum test).

Altogether, these results showed that MB-treated animals behaved like control ones and that the MB treatment in control animals did not alter the physiological somatosensory light and nociceptive thresholds.

Ultimately, to furtherly confirm the postural changes in the diverse experimental conditions, we tracked the paw volar surfaces painted with black ink. Footprints showed comparable shapes in CR (Figure 3D, i), CR-MB (Figure 3D, ii) and CCI-MB (Figure 3D, iv) while strong differences were recoverable in the CP footprints (Figure 3D, iii).

### Electrophysiological experiments

Acute electrophysiology recordings of the somatosensory thalamocortical complex were performed in all four groups of experimental animals (CR, CCI-MB, CCI, CR-MB) by means of couples of microelectrode matrices. Two experimental conditions were considered: the resting state and the stimulated state where precise and fine-tuned tactile stimulations of the affected hind limb where elicited (Antonio G. Zippo et al. 2013; A.G. Zippo et al. 2015, 2016). We then extracted the functional connectivity between the neuronal activity dichotomized by filtering into spiking activity and local field potentials (A.G. Zippo et al. 2013, 2015, 2016). The analysis of the average firing rate including both cortex and thalamus (Figure 4A) during spontaneous activity showed, as expected, a clear separation among groups, significantly increased in CR with respect to CCI (P = 0.029, N = 276, ranksum test) as well as in CCI-MB compared to CCI (P = 0.031, N = 276, ranksum test). Also, the CR-MB animals showed a normal firing rate, slightly higher than CCI (P = 0.015, N = 276, ranksum test). A comparable scenario held for the tactile evoked activity (CR vs CCI: P = 0.018, N = 268; CCI-MB vs CCI: P = 0.011, N = 268; CR-MB vs CCI: P = 0.009, N = 268, ranksum tests). By analyzing the power spectra of LFP (multitaper method) we found that the band of 10-20 Hz was severely reduced during spontaneous activity only in the CCI group (Figure 4B, Kruskal-Wallis test: P < 0.001, N = 552) while the wavelet coefficients related by the tactile stimulation were similar in all four classes (Figure 4C, Kruskal-Wallis test: P < 0.001, N = 552).

**Figure 4.**
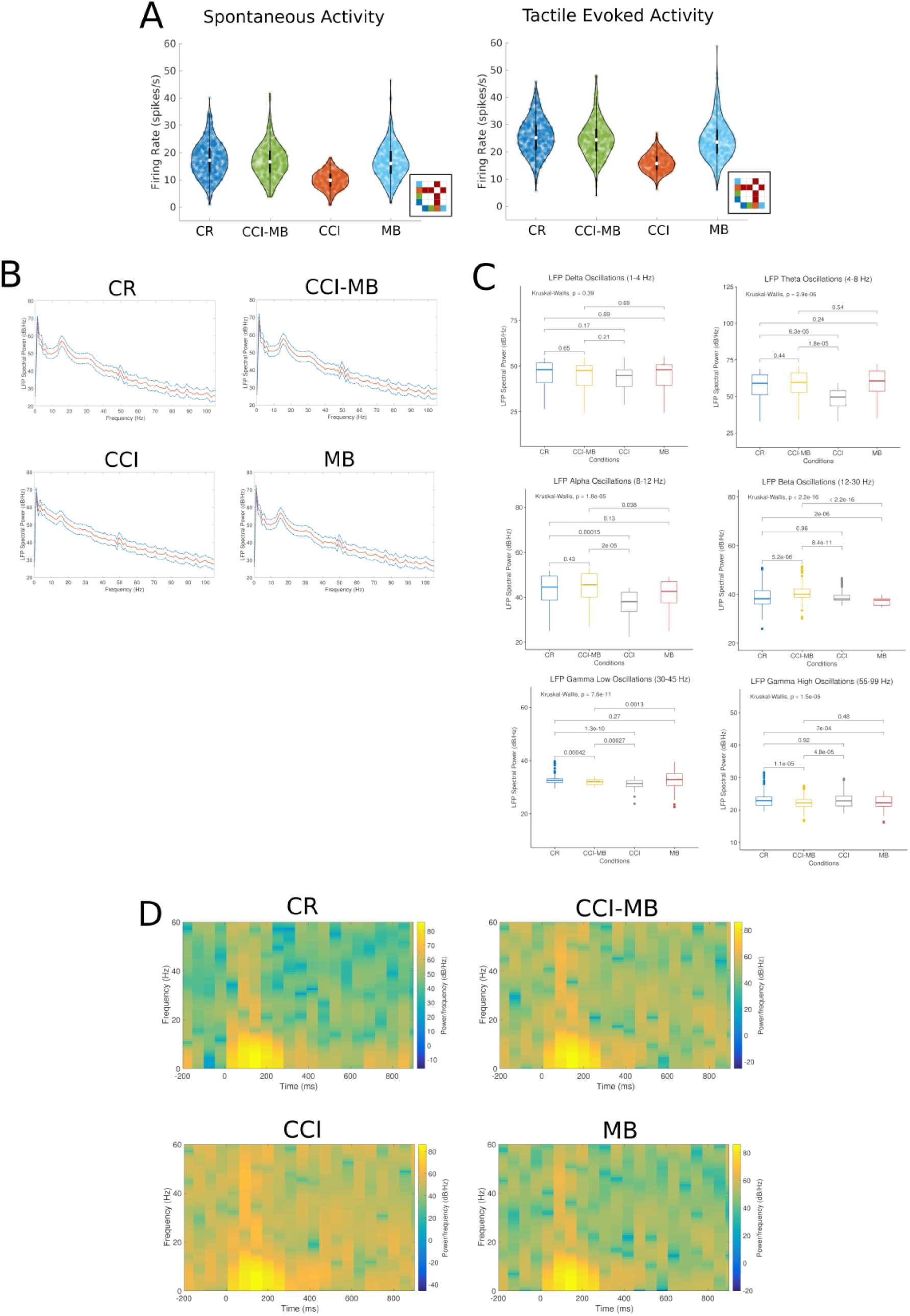
Fundamental electrophysiological results. (A) Firing rate distribution of the recorded neurons both from spontaneous and tactile-evoked activities. The inset matrices indicate the statistical significance (P < 0.05) among groups. Average LFP-power spectra in the four experimental conditions during resting state (B) and in the 6 different LFP bands (C). (D) Average wavelet coefficients of the tactile-evoked activity in the four experimental conditions.

The neuronal functional connections were subsequently analyzed to investigate the mesoscopic scale of neuronal populations. As in the previous works (Antonio G. Zippo et al. 2013; A.G. Zippo et al. 2016), we took into considerations three fundamental complex network statistics, namely the clustering coefficient, the modularity and the efficiency (the inverse of the characteristic path length) in the experimental conditions of resting and tactile evoked states (see Methods) and for spiking and local field potential (LFP) activities. Crucially, we found that the MB treatment restored all network properties lost after the onset of the chronic pain. Specifically, both in the spontaneous (spiking, P < 0.001, N = 276; LFP, P < 0.001, N = 276, ranksum tests) and in the tactile evoked (spiking, P < 0.001, N = 268; LFP, P < 0.001, N = 268, ranksum tests) activities, the loss of efficiency due to the neuropathy (Figure 5A(i)-(ii)) was restored in the MB-treated animals while the MB treatment on control animals (CR-MR) did not provoke any significant modification in comparison to CR animals both on spontaneous (spiking, P = 0.974, N = 276; LFP, P = 0.880, N = 276, ranksum tests) and in evoked activity (spiking, P = 0.811, N = 268; LFP, P = 0.664, N = 268, ranksum tests). Complementarily, the clustering coefficient, tightly reduced in CCI animals in spontaneous (Figure 5(iii)-(iv), spiking, P = 0.001, N = 276; LFP, P = 0.002, N = 276, ranksum tests) and evoked (spiking, P < 0.001, N = 268; LFP, P = 0.009, N = 268, ranksum tests) activity was recovered in the CCI-MB group (spontaneous: spiking, P = 0.944, N = 276; LFP, P = 0.908, N = 276; evoked: spiking, P = 0.623, N = 268; LFP, P = 0.581, N = 268, ranksum tests). Even in this analysis, the MB treatment on control animals did not alter the normal topological properties of the thalamocortical functional networks both in spontaneous (spiking, P = 0.937, N = 276; LFP, P = 0.729, N = 276, ranksum tests) and in evoked (spiking, P = 0.727, N = 268; LFP, P = 0.602, N = 268, ranksum tests) activity.

**Figure 5.**
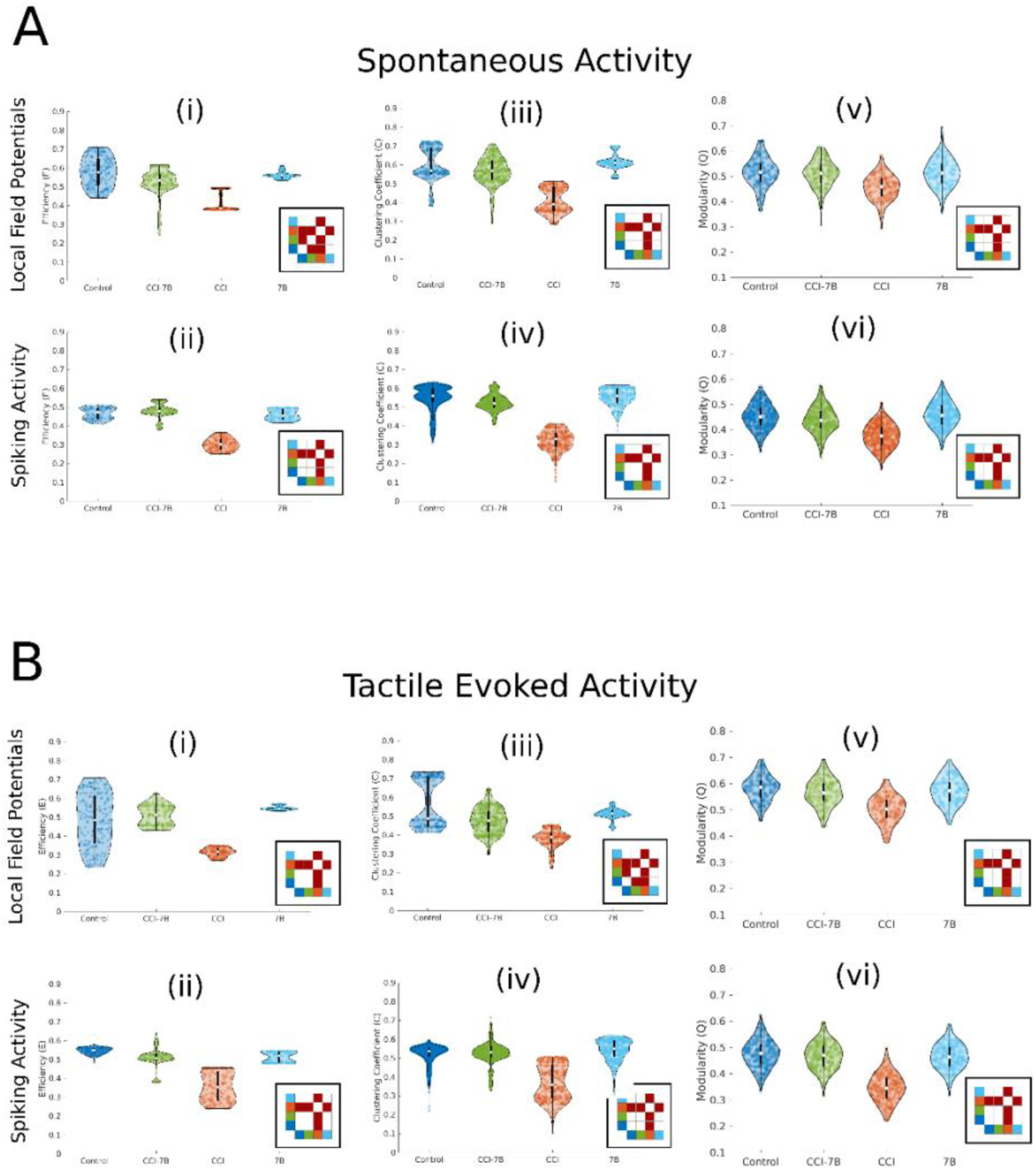
Functional network statistics. (A) Functional connectivity statistics (Efficiency, Clustering Coefficient, Modularity) extracted from the local field potentials (first row, i-iii-v) and spiking activity (second row, ii-iv-vi) in spontaneous state. (B) Functional connectivity statistics extracted from the local field potentials (third row, i-iii-v) and spiking activity (fourth row, ii-iv-vi) in in tactile-evoked state. The inset matrices indicate the statistical significance (P < 0.05) among groups.

At a higher (mesoscopic) scale, the modularity indicated comparable results (Figure 5(v)-(vi)) showing a full recovery in the CCI-MB of the normal modular organization both in spontaneous (spiking, P = 0.770, N = 276; LFP, P = 0.650, N = 276, ranksum tests) and evoked (spiking, P = 0.545, N = 268; LFP, P = 0.877, N = 268, ranksum tests) activity, weakened in the CCI (spontaneous: spiking, P < 0.000, N = 276; LFP, P = 0.003, N = 276; evoked: spiking, P < 0.000, N = 268; LFP, P < 0.000, N = 268, ranksum tests). Analyses showed also that in CR-MB the modularity was not affected by the MBs in spontaneous (spiking, P = 0.652, N = 276; LFP, P = 0.819, N = 276, ranksum tests) and evoked activity (spiking, P = 0.701, N = 268; LFP, P = 0.793, N = 268, ranksum tests).

Eventually, the behavioral and the electrophysiological results confirmed the resolutive efficacy of the MB release to reverse the chronic pain conditions instantiated within the central nervous districts of rat brains.

### SEM analyses

Bone and MB-treated brain scans have been done by Scanning Electron (SEM) and by conventional light transmission microscopies. Acquisitions of the irradiated skull bone have been performed to understand if bone lesions were detectable after the irradiation. No evident traces (20 days after the irradiation, Figure 6A-B) were observable on the bone tiles examined but slight dashes were recoverable (Figure 6C, red dashed lines) assumedly to be related to fast cicatrized MB induced scars. Examinations at higher magnification showed, in fact, traces of scanty active of cell elongations (Figure 6D).

**Figure 6.**
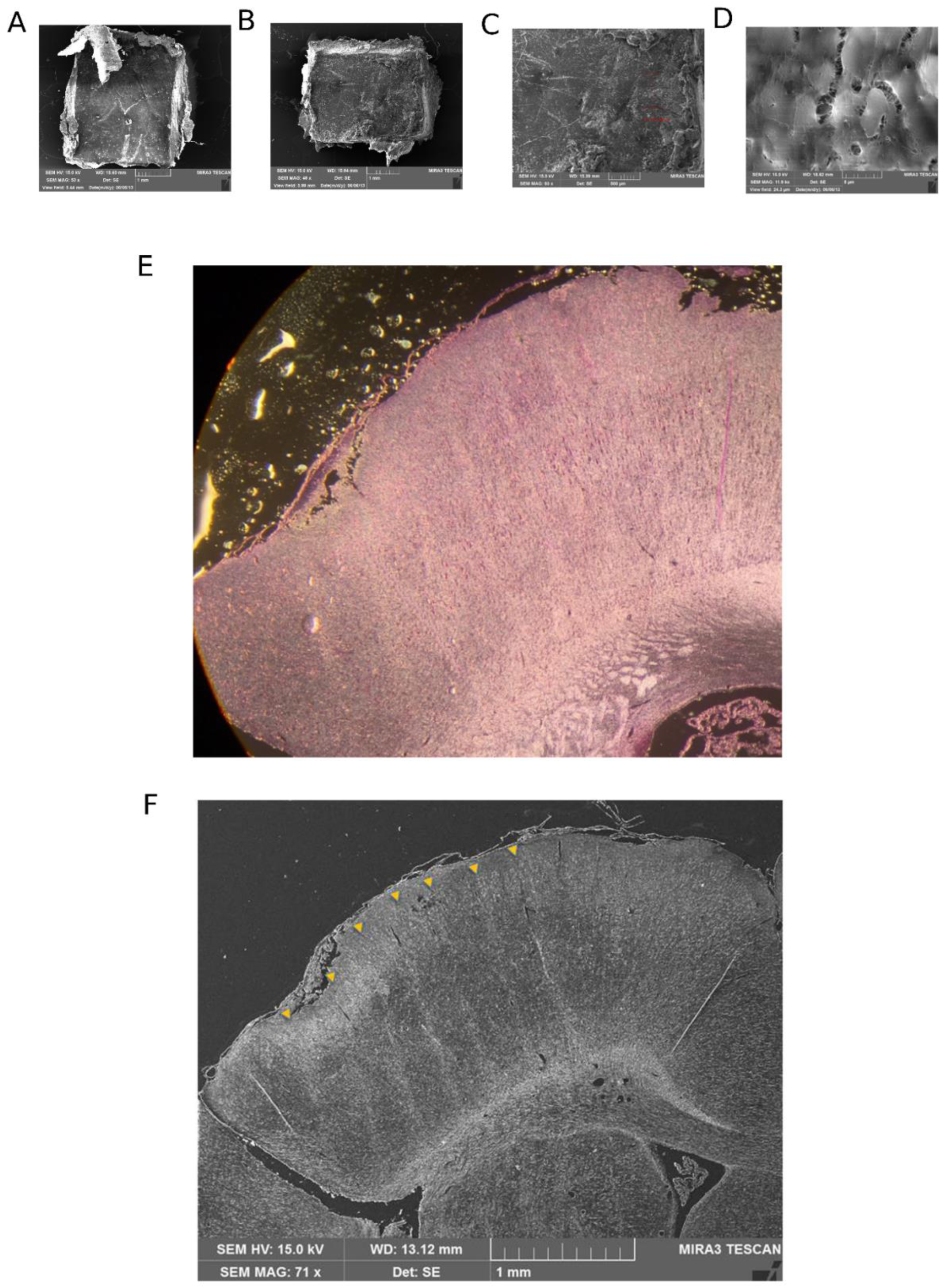
Anatomical confirmations of targeted regions. (A-B) Bone tiles excised from the irradiated skull regions. (C) Higher magnification of a microbeam track in the bone. Red dashed lines indicate the light bone scare trace. (D) Cell-level magnification potentially showing signs of osteoblastic activity in the focus of the beam. (E) Nissl staining and (F) SEM imaging of the X-ray microbeam traces of the coronoal section of the rat brain. The Nissl image is 25 µm behind the slice used in SEM imaging.

Clear tiny traces of the MB irradiation were well evident throughout the whole cortical thickness (Figure 6E). The comparison with the accompanying image in conventional Nissl staining of the same region of the same cortical samples, in a 5 µm thick slice 25 µm posterior to that obtained in SEM imaging along the fronto-occipital axis, confirmed the spatial precision of the MB in their length extension (Figure 6F). These signs appear, in the SEM imaging, significantly fading under the Corpus Callosum region and less easily detectable at the thalamic level. At higher magnification, signs of protein coagulation due to the X-ray MB effects are clearly visible in segregated areas well within the beam tracks (Figure 7B-C). It is interesting to note the neat boundary between the MB involved region and the untouched surrounding tissue. Modest signs of cell elongations appear where the margins of the scar do not exceed the MB width (Figure 7D). This implies that potential secondary degenerations (avalanche degeneration effects) in the tissue are seemingly absent.

**Figure 7.**
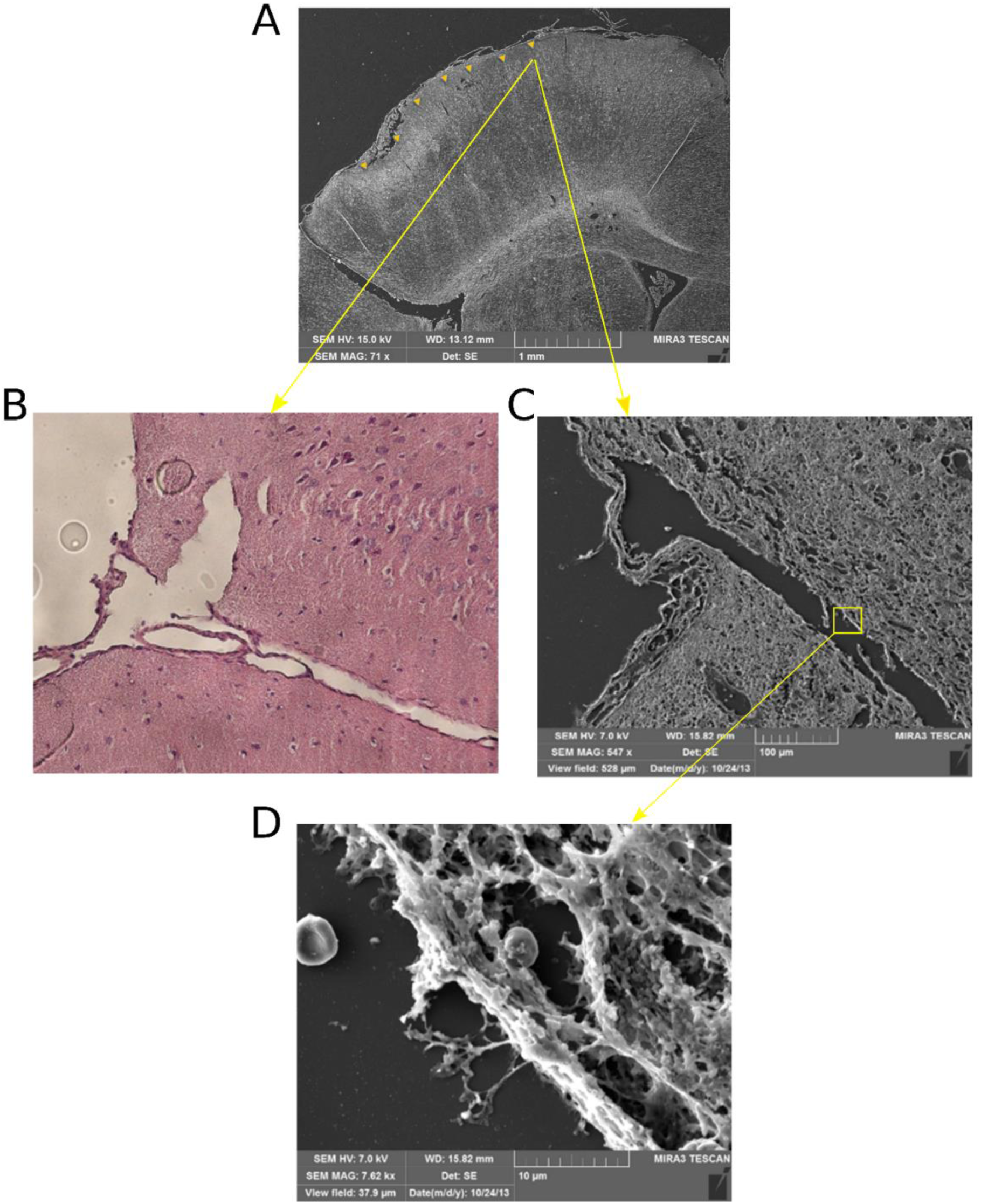
Analysis at progressively higher magnifications of a microbeam track within the cortical tissue. (A) Whole field, (B-C) single beam resolution obtained with Nissl staining and SEM imaging. In (A), microbeams came from the top-left of the figure (the cortical surface) toward the bottom-rigth (deeper brain tissue). The Nissl image is 25 µm behind the slice used in SEM imaging. (D) Further magnification of a detail highlighted in yellow square indicating the coagulation of the microbeam cut border.

Percentage of C, N, O, Na, Mg, Al, Si, Os, S, Ca elements were estimated in all the scans in coherent amounts of the brain tissue, both on direct preparation for SEM and also on slices after treatments for SEM histological examination (Figure 8A, cfr Figure 7C). Direct analyses at higher magnification (Figure 8B, cfr Figure 7D) of beam traces within the brain tissue show slightly higher C levels, potentially due to the X-ray high-dose deposition.

**Figure 8.**
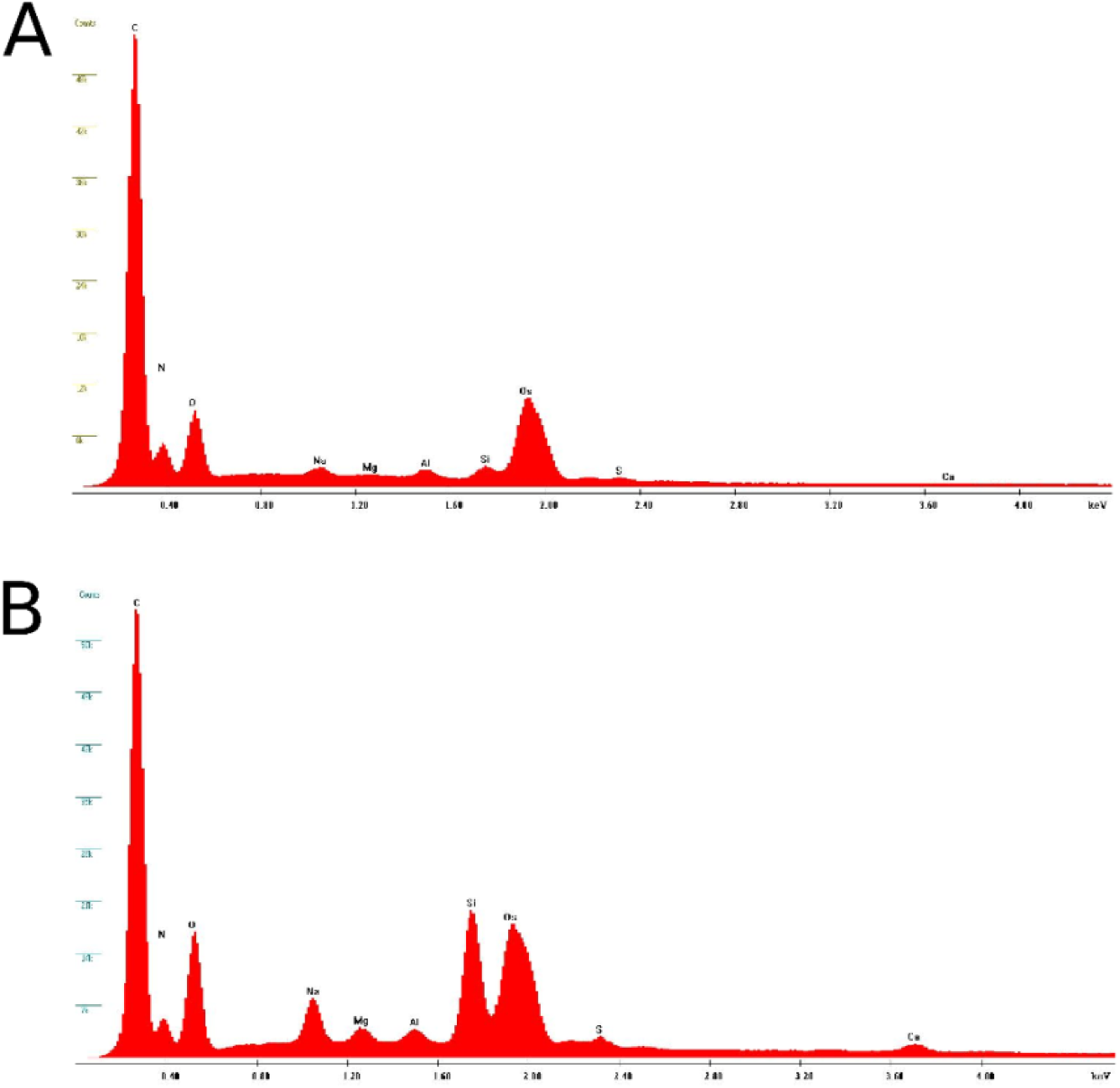
Spectrograms of the composition of the cortical intact tissue (A) and (B) in the region of microbeam cut (cfr. Figure 6D). To note the osmium component of the spectrograms due the SEM preparation, and the silicium high component in the region of cut due to the presence of the slide glass and the higher carbonium component potential congruous with the effects of high-dose X-ray MB depositions.

## Discussion

In this work, we showed that *in vivo* high-dose X-ray microbeam transcranial irradiation of the hind-limb somatosensory cortex in CCI rats abolishes all the behavioral and electrophysiological signs of CP with no detrimental effects when delivered on control animals. Spatially fractionated synchrotron-generated X-rays (with doses of 360 Grey) in form of arrays of seven quasi-parallel, lamellar microbeams, 100 µm wide, and spaced 400 µm center-to-center, were delivered to a set of rats with and without a neuropathic chronic pain model. Local cuts are effective on groups of horizontal neuronal connections (fibers) with these parameters. The ablation of neurons and glial cells along the beam irradiation path becomes negligible in comparison with the efficiency of the functional segregation achieved by the modularization. In fact, the volume of the transected horizontal fibers by the sevenfold microcuts represents a tiny amount of the global cortical connections. However, it efficiently interferes on the local connectivity of the hind-limb sensory cortex fibers leaving seemingly intact all the other sensory contingents. In addition, the MB-resection scars are confined within their borders and non-noticeable in the tissue beyond, excluding microscopically observable lateral degenerative avalanche phenomena outside the beam track path (Studer et al. 2015; Pouyatos et al. 2016; Romanelli et al. 2013).

Diversely from the rich connectivity of the natural cortical (mini)columns, the artificial cortical moduli are effectively mutually isolated by the MB cuts, being however able to communicate via shared thalamo-cortico-thalamic paths (Boucsein et al. 2011; Schnepel et al. 2015; Sherman 2007). Despite these minor sources of mutual communication, we assumed that each artificial module might develop specific own dynamic features after network rearrangements albeit the negligible resections of contingent fibers and neurons (Ellefsen, Mouret, and Clune 2015; Clune et al. 2013; Mengistu et al. 2016). The resultant autonomous anatomical/functional and neuroelectric routes in each independent MB-module, when taken together express a global behavior functionally more proficient than the integer and integral CCI condition. Indeed, this assumption is justified by the seemingly paradoxical increase of information transfer and network graph topological restoration, that we concurrently estimated, and, in addition, by the conversions of pathological sensory anomalies (e.g. hyperpathy, hyperesthesia and hyperalgesia) towards control-like conditions. This goes with the observation that, in biological contexts, modular structures appear more responsive to incoming perturbations, a natural condition in a sensory context (Sporns 2013).

An early support of the work rationale was provided by the *in silico* model that, although the relevant limitation of the anatomo-functional substrate, reproduced the intimate recovering dynamics. Therefore, the ingredients of the reductive model were sufficient for the emergence of the observed phenomena.

Though the broad outcome of the “novel” modular cortex appears mostly realized by rules of functional addition of its modules, this piecewise condition seems anyway adequate to enrich the local information contents to levels matching “normal average” criteria, comparable to control animals (Rowland and Moser 2014). To note that in control animals the lack of MB secondary effects might be ascribed to network resilience due to uncompromised bottom-up and top-down arches of the thalamo-cortico-thalamic loop. A further reason has to be imputed to the surprisingly low sensitivity of normal brain tissue to irradiation (Bräuer-Krisch et al. 2010). If the comparable damage in CCI-MB regions achieves the result to reinstate functionally natural-like conditions, this may suggest that different outcomes could be envisaged depending on the original conditions. Namely, a dynamic shift from a CP signaling by strong horizontal connectivity may undergo a rebalancing by its weakening (Brodal 2007) as confirmed in literature (Schnepel et al. 2015; Angelucci and Bressloff 2006; Négyessy et al. 2013). Indeed, because intrinsic horizontal connections appear crucially involved in the rat primary somatosensory cortex tasks (Négyessy et al. 2013), we can reasonably assume a similar involvement in the pain processing (B A Johnson and Frostig 2016; Brett A Johnson and Frostig 2018). The advantage of a conversion from a CP statistically stationary structure to an artificial modular complex able to augment and improve the information content is ratified, also, by the potential advantages of partitioned architectures, as shown in many biological contexts (Bullmore and Sporns 2012). The specific choice to generate moduli is, in fact, related to the idea of natural Brain Modularity, a key driver for evolving nervous systems, granting for better adaptability (Ellefsen, Mouret, and Clune 2015; Clune et al. 2013). In evolution, Modularity appears a suitable answer to selection pressures to maximize network performance and minimize connection costs (Clune et al. 2013; Gómez-Robles, Hopkins, and Sherwood 2014). Besides these functional connectivity issues of CP, correlated problems such as the increased energy costs of the involved networks seem to become more manageable. Modularization may also help or concur, indeed, to reduce a global somatosensory metabolic problem into smaller and more manageable contexts (Bullmore and Sporns 2012). In confirming these findings, the topological approach by graph theory to network architectures helps to clarify and potentially deepen these issues. After the X-ray microbeam delivery, the reduced information in CP, mirrored in the graph topology degradation from *modules-and-hubs* to random architectures, shows a novel conversion to control-like topology (A.G. Zippo et al. 2016). Behaviorally, this conversion appears reflected in the reoccurrence of normal-like features, suggesting that sensory parameters may be either revived or *ex-novo* regenerated by the cortical X-ray manipulation. In the former case, specific parameters such as the many plasticity events continuously occurring in the normal somatosensory cortex may revive in the presence of strongly modulating events. Redundant cortical functions may be thus more efficient in the closed context of moduli, as in control animals where the network redundancy is resilient to the low grade degradation from MB (Gao, Barzel, and Barabási 2016). The alternative hypothesis of regeneration of properties appears weaker, in that no specific neurogenesis after irradiation has been observed in the literature (Brönnimann et al. 2016) (except for poor potential cell extensions as shown in Figure 7). The irradiation induces also local vascular regenerative episodes where signs of novel vessel branches transpassing narrower MB tracks (25 µm, 500 Gy) have been observed (Serduc et al. 2008).

On the electrophysiological side, the EEG recordings taken from 4 leads during the experiments did not show significant power spectral differences among the four conditions (data not shown, see (A.G. Zippo et al. 2016)). At the neuronal level, the recovery of the network topology significantly increased towards levels comparable to controls. Specifically, the spiking and the Local Field Potential power spectra that, grossly represent the unit and the synaptic dynamics, in CCI-MB rats *per se* re-presented statistical control-like parameters. This appears to emphasize the importance of the local intramodular connectome in respect of a global dynamics and that moduli formation could generate neurodynamic contexts more susceptible to recover collapsed features (Ellefsen, Mouret, and Clune 2015). From a network perspective, though the vertical connectivity has largely been explored within the cortical layers (Thomson and Bannister 2003; Levy and Reyes 2012), the horizontal connections (both parallel for homolaminar and oblique for interlaminar connections) are gaining importance. This is true in many cortical operations such as the diffusion of signals within sensory areas (Mao et al. 2011), in surround suppression (the weakening of fringe excitations enhancing the stimulus detection (Adesnik et al. 2012)) or in complex mechanisms of multiple contextual inputs coherence (Petreanu et al. 2012). Recently, a hypothesis has been advanced that the majority of synaptic connections (∼ 75%) a cortical neuron receives, actually originates from outside the local volume (Stepanyants et al. 2009; Boucsein et al. 2011), even dominating the cortical network dynamics (Schnepel et al. 2015). Despite the potential synaptic strength differences, horizontal connections seem thus to promote robust asynchronous ongoing activity states and reduce noise correlations in stimulus-induced activity (Boucsein et al. 2011). It is possible to hypothesize a double gating that, in presence of a horizontal connectivity efficacy excess, a pruning operation could restitute a sound synaptic strength to the cortical circuitry. Another issue to be weighted relates to the depth of the X-ray penetration that goes far beyond the cortical mantle towards the subcortical structures (Romanelli et al. 2013; Serduc et al. 2008; Studer et al. 2015; Pouyatos et al. 2016). This means that, though weakened in their path through the cerebral matter, some degree of damage could affect the tissues beneath the cerebral cortex. Most of the connections are indeed intact and not directly involved by the X-ray beams. As from the literature (Studer et al. 2015), however, with the used irradiation parameters no gross damage to the neural structure should be evident anyway.

Along these positive theoretical and empirical results, the experimental setting enlightens also a proof-of-concept and seems a suitable model for potential translational medicine with some important caveats, first of all taking into account the human gyrencephalic cortex in contrast with the rodent lissencephalic cortex. A hypothetical translation could benefit of the multiangular, modulated X-ray beam strategy to selectively target the complex geometries of the neural structures of interest. Eventually, as a whole, the experimental results further indicate that chronic pain may be deeply reconsidered in the light of connectivity disorders at the central level and that suitable treatments could be envisaged in complex practices apparently very far from the conventional ones.

## Author contributions

AGZ performed all the analyses, wrote the paper, GB performed the histological examinations, MPR did the SEM imaging, MV performed the behavioral and the electrophysiological experiments, HR prepared and performed the Synchrotron beamline irradiation, GCC did the technical mechatronic control of the neuronal recordings, EBK performed the dosimetry and estimated the peak-valley X-ray MB Doses, VDG performed some data analyses and helped in the CP model generation, PC planned the synchrotron parameters and guided the image analysis, AB planned the Synchrotron analyses, performed the Synchrotron irradiations, directed the ESRF Synchrotron personnel during the experiments, GEMB planned the whole experimental cycle, performed the experiments (Synchrotron, SEM and Electrophysiology), wrote the paper.

### Acknowledgments

The biomedical facility of the ESRF is acknowledged for the support in the management and care of animals. Authors acknowledge the support provided by the TD1205 COST Action. The authors are grateful to Mr. Giacomo Barbone, PhD student at the Ludwig-Maximilians-Universität (München, Deutschland), for the careful English proof-reading.

---

**Table 1.**
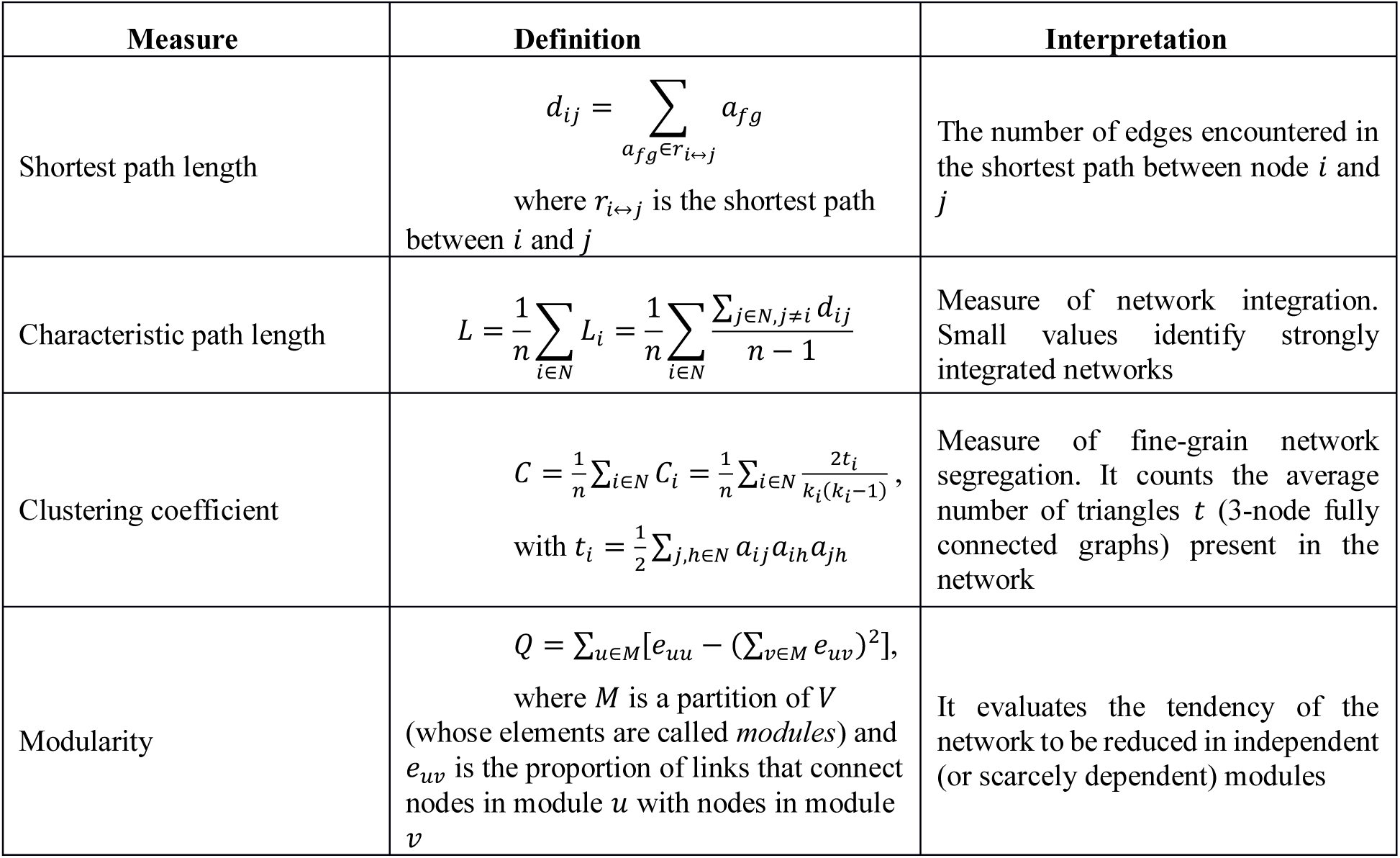
Complex network statistics commonly used in brain connectomics. All formulas are referred to a (undirected) graph 〈*V*, *E*〉, with |*V*| = *N*, opportunely described by the adjacency *N*×*N*-matrix *A*= *a*_*ij*_ where *a*_*ij*_=1 if and only if there exist the element (*i,j*) in the set *E* and 0 otherwise.

| Measure | Definition | Interpretation |
| --- | --- | --- |
| Shortest path length | $d_{ij} = \sum_{a_{fg} \in r_{i \leftrightarrow j}} a_{fg}$<br>where $r_{i \leftrightarrow j}$ is the shortest path between $i$ and $j$ | The number of edges encountered in the shortest path between node $i$ and $j$ |
| Characteristic path length | $L = \frac{1}{n} \sum_{i \in N} L_i = \frac{1}{n} \sum_{i \in N} \frac{\sum_{j \in N, j \neq i} d_{ij}}{n-1}$ | Measure of network integration. Small values identify strongly integrated networks |
| Clustering coefficient | $C = \frac{1}{n} \sum_{i \in N} C_i = \frac{1}{n} \sum_{i \in N} \frac{2t_i}{k_i(k_i-1)},$<br>with $t_i = \frac{1}{2} \sum_{j, h \in N} a_{ij} a_{ih} a_{jh}$ | Measure of fine-grain network segregation. It counts the average number of triangles $t$ (3-node fully connected graphs) present in the network |
| Modularity | $Q = \sum_{u \in M} [e_{uu} - (\sum_{v \in M} e_{uv})^2],$<br>where $M$ is a partition of $V$ (whose elements are called *modules*) and $e_{uv}$ is the proportion of links that connect nodes in module $u$ with nodes in module $v$ | It evaluates the tendency of the network to be reduced in independent (or scarcely dependent) modules |

**Movies 1-4**. Samples of walking patterns in CR (1), CCI (2), CCI-MB (3), CR-MB (4) animals.

## References

Adesnik, Hillel, William Bruns, Hiroki Taniguchi, Z. Josh Huang, and Massimo Scanziani. 2012. “A Neural Circuit for Spatial Summation in Visual Cortex.” Nature 490 (7419): 226–31. https://doi.org/10.1038/nature11526.

Angelucci, Alessandra, and Paul C. Bressloff. 2006. “Contribution of Feedforward, Lateral and Feedback Connections to the Classical Receptive Field Center and Extra-Classical Receptive Field Surround of Primate V1 Neurons.” Progress in Brain Research 154: 93–120. https://doi.org/10.1016/S0079-6123(06)54005-1.

Apkarian, A, M N Baliki, and Paul Y Geha. 2009. “Towards a Theory of Chronic Pain.” Progress in Neurobiology 87 (2). Pergamon: 81–97. https://doi.org/10.1016/J.PNEUROBIO.2008.09.018.

Babadi, Baktash, Haim Sompolinsky, K. Bhaukaurally, J. Beroud, A. Carleton, M. Mendelsohn, J. Edmondson, R. Axel, F.E. Hoebeek, and C.I. De Zeeuw. 2014. “Sparseness and Expansion in Sensory Representations.” Neuron 83 (5). Neural Information Processing Systems, La Jolla, CA: 1213–26. https://doi.org/10.1016/j.neuron.2014.07.035.

Basbaum, Allan I, Diana M Bautista, Gregory Scherrer, and David Julius. 2009. “Cellular and Molecular Mechanisms of Pain.” Cell 139 (2). United States: 267–84. https://doi.org/10.1016/j.cell.2009.09.028.

Bennett, G J, and Y K Xie. 1988. “A Peripheral Mononeuropathy in Rat That Produces Disorders of Pain Sensation like Those Seen in Man.” Pain 33 (1). NETHERLANDS: 87–107.

Boucsein, Clemens, Martin Nawrot, Philipp Schnepel, and Ad Aertsen. 2011. “Beyond the Cortical Column: Abundance and Physiology of Horizontal Connections Imply a Strong Role for Inputs from the Surround.” Frontiers in Neuroscience 5. Frontiers: 32. https://doi.org/10.3389/fnins.2011.00032.

Bräuer-Krisch, E., H. Requardt, T. Brochard, G. Berruyer, M. Renier, J. A. Laissue, and A. Bravin. 2009. “New Technology Enables High Precision Multislit Collimators for Microbeam Radiation Therapy.” Review of Scientific Instruments 80 (7): 074301. https://doi.org/10.1063/1.3170035.

Bräuer-Krisch, E., R. Serduc, E.A. Siegbahn, G. Le Duc, Y. Prezado, A. Bravin, H. Blattmann, and J.A. Laissue. 2010. “Effects of Pulsed, Spatially Fractionated, Microscopic Synchrotron X-Ray Beams on Normal and Tumoral Brain Tissue.” Mutation Research/Reviews in Mutation Research 704 (1–3). Elsevier: 160–66. https://doi.org/10.1016/J.MRREV.2009.12.003.

Brodal, Per. 2007. “The Central Nervous System.” In, 490–93. Oxford Univ. Press.

Brönnimann, Daniel, Audrey Bouchet, Christoph Schneider, Marine Potez, Raphaël Serduc, Elke Bräuer-Krisch, Werner Graber, Stephan von Gunten, Jean Albert Laissue, and Valentin Djonov. 2016. “Synchrotron Microbeam Irradiation Induces Neutrophil Infiltration, Thrombocyte Attachment and 8 Selective Vascular Damage in Vivo.” Scientific Reports 6 (1). Nature Publishing Group: 33601. https://doi.org/10.1038/srep33601.

Bullmore, Ed, and Olaf Sporns. 2012. “The Economy of Brain Network Organization.” Nature Reviews. Neuroscience 13 (5): 336–49. https://doi.org/10.1038/nrn3214.

Clune, Jeff, Jean-baptiste Mouret, Hod Lipson, Jeff Clune, Jean-baptiste Mouret, and Hod Lipson. 2013. “The Evolutionary Origins of Modularity The Evolutionary Origins of Modularity,” no. January.

Coan, Paola, Angela Peterzol, Stefan Fiedler, Cyril Ponchut, Jean Claude Labiche, and Alberto Bravin. 2006. “Evaluation of Imaging Performance of a Taper Optics CCD ‘FReLoN’ Camera Designed for Medical Imaging.” Journal of Synchrotron Radiation 13 (3): 260–70. https://doi.org/10.1107/S0909049506008983.

Crosbie, Jeffrey C., Pauline Fournier, Stefan Bartzsch, Mattia Donzelli, Iwan Cornelius, Andrew W. Stevenson, Herwig Requardt, and Elke Bräuer-Krisch. 2015. “Energy Spectra Considerations for Synchrotron Radiotherapy Trials on the ID17 Bio-Medical Beamline at the European Synchrotron Radiation Facility.” Journal of Synchrotron Radiation 22 (4): 1035–41. https://doi.org/10.1107/S1600577515008115.

Darian-Smith, Corinna, and Charles D. Gilbert. 1994. “Axonal Sprouting Accompanies Functional Reorganization in Adult Cat Striate Cortex.” Nature 368 (6473). Nature Publishing Group: 737–40. https://doi.org/10.1038/368737a0.

Descalzi, Giannina, Daigo Ikegami, Toshikazu Ushijima, Eric J Nestler, Venetia Zachariou, and Minoru Narita. 2015. “Epigenetic Mechanisms of Chronic Pain.” Trends in Neurosciences 38 (4). England: 237–46. https://doi.org/10.1016/j.tins.2015.02.001.

Ellefsen, Kai Olav, Jean-Baptiste Mouret, and Jeff Clune. 2015. “Neural Modularity Helps Organisms Evolve to Learn New Skills without Forgetting Old Skills.” Edited by Josh C. Bongard. PLOS Computational Biology 11 (4). Public Library of Science: e1004128. https://doi.org/10.1371/journal.pcbi.1004128.

Francis, Joseph Thachil, and Weiguo Song. 2011. “Neuroplasticity of the Sensorimotor Cortex during Learning.” Neural Plasticity 2011 (September). Hindawi: 310737. https://doi.org/10.1155/2011/310737.

Gao, Jianxi, Baruch Barzel, and Albert-László Barabási. 2016. “Universal Resilience Patterns in Complex Networks.” Nature 530 (7590). Nature Publishing Group: 307–12. https://doi.org/10.1038/nature16948.

Gårdmark, M, A U Höglund, and M Hammarlund-Udenaes. 1998. “Aspects on Tail-Flick, Hot-Plate and Electrical Stimulation Tests for Morphine Antinociception.” Pharmacology & Toxicology 83 (6): 252–58. http://www.ncbi.nlm.nih.gov/pubmed/9868743.

Gómez-Robles, Aida, William D Hopkins, and Chet C Sherwood. 2014. “Modular Structure Facilitates Mosaic Evolution of the Brain in Chimpanzees and Humans.” Nature Communications 5 (July). NIH Public Access: 4469. https://doi.org/10.1038/ncomms5469.

Gustin, Sylvia M, Chris C Peck, Lukas B Cheney, Paul M Macey, Greg M Murray, and Luke A Henderson. 2012. “Development/Plasticity/Repair Pain and Plasticity: Is Chronic Pain Always Associated with Somatosensory Cortex Activity and Reorganization?” The Journal of Neuroscience 32 (43): 14874–84. https://doi.org/10.1523/JNEUROSCI.1733-12.2012.

Johnson, B A, and R D Frostig. 2016. “Long, Intrinsic Horizontal Axons Radiating through and beyond Rat Barrel Cortex Have Spatial Distributions Similar to Horizontal Spreads of Activity Evoked by Whisker Stimulation.” Brain Structure & Function 221 (7). NIH Public Access: 3617–39. https://doi.org/10.1007/s00429-015-1123-7.

Johnson, Brett A, and Ron D Frostig. 2018. “Long-Range, Border-Crossing, Horizontal Axon Radiations Are a Common Feature of Rat Neocortical Regions That Differ in Cytoarchitecture.” Frontiers in Neuroanatomy 12. Frontiers Media SA: 50. https://doi.org/10.3389/fnana.2018.00050.

Laird, J M, and G J Bennett. 1993. “An Electrophysiological Study of Dorsal Horn Neurons in the Spinal Cord of Rats with an Experimental Peripheral Neuropathy.” Journal of Neurophysiology 69 (6). UNITED STATES: 2072–85.

Levy, R. B., and A. D. Reyes. 2012. “Spatial Profile of Excitatory and Inhibitory Synaptic Connectivity in Mouse Primary Auditory Cortex.” Journal of Neuroscience 32 (16): 5609–19. https://doi.org/10.1523/JNEUROSCI.5158-11.2012.

Liang, Lingli, Brianna Marie Lutz, Alex Bekker, and Yuan-Xiang Tao. 2015. “Epigenetic Regulation of Chronic 9 Pain.” Epigenomics 7 (2). England: 235–45. https://doi.org/10.2217/epi.14.75.

Mansour, A R, M A Farmer, M N Baliki, and A Vania Apkarian. 2014. “Chronic Pain: The Role of Learning and Brain Plasticity.” Restorative Neurology and Neuroscience 32 (1). NIH Public Access: 129–39. https://doi.org/10.3233/RNN-139003.

Mao, Tianyi, Deniz Kusefoglu, Bryan M Hooks, Daniel Huber, Leopoldo Petreanu, and Karel Svoboda. 2011. “Long-Range Neuronal Circuits Underlying the Interaction between Sensory and Motor Cortex.” Neuron 72 (1): 111–23. https://doi.org/10.1016/j.neuron.2011.07.029.

Melzack, R. 2001. “Pain and the Neuromatrix in the Brain.” Journal of Dental Education 65 (12): 1378–82. http://www.ncbi.nlm.nih.gov/pubmed/11780656.

Mengistu, Henok, Joost Huizinga, Jean-Baptiste Mouret, and Jeff Clune. 2016. “The Evolutionary Origins of Hierarchy.” Edited by Olaf Sporns. PLOS Computational Biology 12 (6). Public Library of Science: e1004829. https://doi.org/10.1371/journal.pcbi.1004829.

Millan, M J, A Członkowski, C W Pilcher, O F Almeida, M H Millan, F C Colpaert, and A Herz. 1987. “A Model of Chronic Pain in the Rat: Functional Correlates of Alterations in the Activity of Opioid Systems.” The Journal of Neuroscience: The Official Journal of the Society for Neuroscience 7 (1): 77–87. http://www.ncbi.nlm.nih.gov/pubmed/3027278.

Narula, Vaibhav, Antonio Giuliano Zippo, Alessandro Muscoloni, Gabriele Eliseo M. Biella, and Carlo Vittorio Cannistraci. 2017. “Can Local-Community-Paradigm and Epitopological Learning Enhance Our Understanding of How Local Brain Connectivity Is Able to Process, Learn and Memorize Chronic Pain?” Applied Network Science 2 (1). Springer International Publishing: 28. https://doi.org/10.1007/s41109-017-0048-x.

Négyessy, László, Emese Pálfi, Mária Ashaber, Cory Palmer, Balázs Jákli, Robert M. Friedman, Li M. Chen, and Anna W. Roe. 2013. “Intrinsic Horizontal Connections Process Global Tactile Features in the Primary Somatosensory Cortex: Neuroanatomical Evidence.” Journal of Comparative Neurology 521 (12): 2798–2817. https://doi.org/10.1002/cne.23317.

Paxinos, G, and C Watson. 1998. “The Rat Brain Atlas in Stereotaxic Coordinates.” San Diego: Academic.

Petreanu, Leopoldo, Diego A. Gutnisky, Daniel Huber, Ning-long Xu, Dan H. O’Connor, Lin Tian, Loren Looger, and Karel Svoboda. 2012. “Activity in Motor–Sensory Projections Reveals Distributed Coding in Somatosensation.” Nature 489 (7415): 299–303. https://doi.org/10.1038/nature11321.

Pouyatos, B, C Nemoz, T Chabrol, M Potez, E Bräuer, L Renaud, K Pernet-Gallay, et al. 2016. “Synchrotron X-Ray Microtransections: A Non Invasive Approach for Epileptic Seizures Arising from Eloquent Cortical Areas.” Scientific Reports 6 (1): 27250. https://doi.org/10.1038/srep27250.

Price, Theodore J., Allan I. Basbaum, Jacqueline Bresnahan, Jan F. Chambers, Yves De Koninck, Robert R. Edwards, Ru-Rong Ji, et al. 2018. “Transition to Chronic Pain: Opportunities for Novel Therapeutics.” Nature Reviews Neuroscience, May. Nature Publishing Group, 1. https://doi.org/10.1038/s41583-018-0012-5.

Romanelli, Pantaleo, Erminia Fardone, Giuseppe Battaglia, Elke Bräuer-Krisch, Yolanda Prezado, Herwig Requardt, Geraldine Le Duc, et al. 2013. “Synchrotron-Generated Microbeam Sensorimotor Cortex Transections Induce Seizure Control without Disruption of Neurological Functions.” Edited by Michael Lim. PloS One 8 (1): e53549. https://doi.org/10.1371/journal.pone.0053549.

Rowland, David C, and May-Britt Moser. 2014. “From Cortical Modules to Memories.” Current Opinion in Neurobiology 24: 22–27. https://doi.org/10.1016/j.conb.2013.08.012.

Schnepel, Philipp, Arvind Kumar, Mihael Zohar, Ad Aertsen, and Clemens Boucsein. 2015. “Physiology and Impact of Horizontal Connections in Rat Neocortex.” Cerebral Cortex 25 (10): 3818–35. https://doi.org/10.1093/cercor/bhu265.

Serduc, Raphaël, Thomas Christen, Jean Laissue, Régine Farion, Audrey Bouchet, Boudewijn van der Sanden, Christoph Segebarth, et al. 2008. “Brain Tumor Vessel Response to Synchrotron Microbeam Radiation Therapy: A Short-Term in Vivo Study.” Physics in Medicine and Biology 53 (13): 3609–22. https://doi.org/10.1088/0031-9155/53/13/015.

Sherman, S Murray. 2007. “The Thalamus Is More than Just a Relay.” Current Opinion in Neurobiology 17 (4). NIH Public Access: 417–22. https://doi.org/10.1016/j.conb.2007.07.003.

Siegbahn, E.A., E. Bräuer-Krisch, J. Stepanek, H. Blattmann, J.A. Laissue, and A. Bravin. 2005. “Dosimetric Studies of Microbeam Radiation Therapy (MRT) with Monte Carlo Simulations.” Nuclear Instruments and Methods in Physics Research Section A: Accelerators, Spectrometers, Detectors and Associated Equipment 548 (1): 54–58. https://doi.org/10.1016/j.nima.2005.03.065.

Spisák, Tamás, Zsófia Pozsgay, Csaba Aranyi, Szabolcs Dávid, Pál Kocsis, Gabriella Nyitrai, Dávid Gajári, Miklós Emri, András Czurkó, and Zsigmond Tamás Kincses. 2017. “Central Sensitization-Related Changes of Effective and Functional Connectivity in the Rat Inflammatory Trigeminal Pain Model.” Neuroscience 344 (March): 133–47. https://doi.org/10.1016/j.neuroscience.2016.12.018.

Sporns, Olaf. 2013. “Structure and Function of Complex Brain Networks.” Dialogues in Clinical Neuroscience 15 (3): 247–62. http://www.ncbi.nlm.nih.gov/pubmed/24174898.

Stepanyants, A., L. M. Martinez, A. S. Ferecsko, and Z. F. Kisvarday. 2009. “The Fractions of Short- and Long-Range Connections in the Visual Cortex.” Proceedings of the National Academy of Sciences 106 (9): 3555–60. https://doi.org/10.1073/pnas.0810390106.

Studer, F., R. Serduc, B. Pouyatos, T. Chabrol, E. Bräuer-Krisch, M. Donzelli, C. Nemoz, J.A. Laissue, F. Estève, and A. Depaulis. 2015. “Synchrotron X-Ray Microbeams: A Promising Tool for Drug-Resistant Epilepsy Treatment.” Physica Medica 31 (6): 607–14. https://doi.org/10.1016/j.ejmp.2015.04.005.

Thomson, Alex M, and A Peter Bannister. 2003. “Interlaminar Connections in the Neocortex.” Cerebral Cortex (New York, N.Y.: 1991) 13 (1): 5–14. http://www.ncbi.nlm.nih.gov/pubmed/12466210.

Tomsett, Richard J., Matt Ainsworth, Alexander Thiele, Mehdi Sanayei, Xing Chen, Marc A. Gieselmann, Miles A. Whittington, Mark O. Cunningham, and Marcus Kaiser. 2015. “Virtual Electrode Recording Tool for EXtracellular Potentials (VERTEX): Comparing Multi-Electrode Recordings from Simulated and Biological Mammalian Cortical Tissue.” Brain Structure and Function 220 (4). Springer Berlin Heidelberg: 2333–53. https://doi.org/10.1007/s00429-014-0793-x.

Vierck, Charles J, Barry L Whitsel, Oleg V Favorov, Alexander W Brown, and Mark Tommerdahl. 2013. “Role of Primary Somatosensory Cortex in the Coding of Pain.” Pain 154 (3). NIH Public Access: 334–44. https://doi.org/10.1016/j.pain.2012.10.021.

Ziegler, E. A., W. Magerl, R. A. Meyer, and R. D. Treede. 1999. “Secondary Hyperalgesia to Punctate Mechanical Stimuli: Central Sensitization to A-Fibre Nociceptor Input.” Brain 122 (12): 2245–2257.

Zippo, A.G., and I. Castiglioni. 2016. “Integration of^18^FDG-PET Metabolic and Functional Connectomes in the Early Diagnosis and Prognosis of the Alzheimer’s Disease.” Current Alzheimer Research 13 (5).

Zippo, A.G., S. Nencini, G.C. Caramenti, M. Valente, R. Storchi, and G.E.M. Biella. 2014. “A Simple Stimulatory Device for Evoking Point-like Tactile Stimuli: A Searchlight for LFP to Spike Transitions.” Journal of Visualized Experiments, no. 85. https://doi.org/10.3791/50941.

Zippo, A.G., S. Rinaldi, G. Pellegata, G.C. Caramenti, M. Valente, V. Fontani, and G.E.M. Biella. 2015. “Electrophysiological Effects of Non-Invasive Radio Electric Asymmetric Conveyor (REAC) on Thalamocortical Neural Activities and Perturbed Experimental Conditions.” Scientific Reports 5. https://doi.org/10.1038/srep18200.

Zippo, A.G., R. Storchi, S. Nencini, G.C. Caramenti, M. Valente, and G.M. Biella. 2013. “Neuronal Functional Connection Graphs among Multiple Areas of the Rat Somatosensory System during Spontaneous and Evoked Activities.” PLoS Computational Biology 9 (6). https://doi.org/10.1371/journal.pcbi.1003104.

Zippo, A.G., M. Valente, G.C. Caramenti, and G.E.M. Biella. 2016. “The Thalamo-Cortical Complex Network Correlates of Chronic Pain.” Scientific Reports 6. https://doi.org/10.1038/srep34763.

Zippo, Antonio G., Riccardo Storchi, Sara Nencini, Gian Carlo Caramenti, Maurizio Valente, and Gabriele Eliseo M. Biella. 2013. “Neuronal Functional Connection Graphs among Multiple Areas of the Rat Somatosensory System during Spontaneous and Evoked Activities.” Edited by Olaf Sporns. PLoS Computational Biology 9 (6). United States: Public Library of Science: e1003104. https://doi.org/10.1371/journal.pcbi.1003104.

