## Supplementary Materials and Methods for "Removal of behavioural and electrophysiological signs of chronic pain by *in vivo* microsections of rat somatosensory cortex with parallel X-ray microbeams"

[†]this work is in memory of Dr. Elke Bräuer-Krisch

### Materials and Methods

#### Animal Surgery for the neuropathic Chronic Constriction Injury (CCI or Bennett-Xie model)

28 animals have been used, 7 for each experimental group: Control Animals (CR), Chronic Constriction Injury (CCI or Bennett-Xie model) animals, microbeam treated CCI animals (CCI-MB) and microbeam treated control animals (CR-MB). Animals were 10 weeks old. All the surgical procedures have been already described in (A.G. Zippo et al. 2016, 2015; Bennett and Xie 1988; Narula et al. 2017). Briefly, prior to surgery to generate the neuropathy, all the materials for the nerve constriction and the surgical sutures were prepared. The 4-0 chromic gut suture sterile wire was cut in segments of 10 cm and immersed in Betadine solution first and in sterile saline after. The animal anesthesia was performed by intraperitoneal (i.p.) sodium pentobarbital (50 mg/kg).
After the full development of anesthesia, the fur on the posterior right thigh was shaved and the skin cleaned with Betadine. The rats were then placed on a heating pad, the hind paw positioned at 90° relative to the backbone. A 3- to 4-cm incision was then performed down the center of the posterior thigh (well identifiable by finger starting from the lumbar vertebrae) parallel to the femoral bone.
After the skin incision, the biceps femoris muscle fascicles were delicately separated by blunt-tipped surgical tweezers and the surgical wound lips of the muscle widened until a 1 cm tract of the sciatic nerve could be visualized. Under a dissecting microscope, thin surgical forceps were used to introduce a curved needle mounted with the sterile gut suture wire under the top end of the visualized sciatic nerve tract, around the nerve. Without stretching the nerve, the chromic gut suture wire was then advanced until the needle was completely passing to the other side of the nerve pulling the wire until both side lengths were equal. A ligature around the nerve with a very loose single loop was then performed and the knot loosely tightened, trimming the two wires end at 0.5 mm above the knot. The maneuver was then repeated threefold along the nerve at 1 mm apart from the previous ligature until the down end was reached. A total of four ligatures along the nerve 1 mm each apart was the final result. The muscle wound lips were then closed in layers by absorbable suture wire. Skin suture wire was then used to close the wound. The animals were allowed to recover in a warm, temperature-controlled cage with a solid floor.

#### Synchrotron X-Ray Microbeam Delivery

The rats for the CCI Chronic Pain model were randomly chosen from the group of rats selected for the experiments (28 rats, 7 rats for experimental group as shown above). The 7 CCI-MB rats were MB treated 15-18 days after the CCI chronic pain model generation. Rats undergoing CCI model generation with no MB treatment have equally been randomly chosen from the original group of rats. Half of control rats (7 over 14) undergoing MB delivery were equally randomly selected.

*X-ray microbeams source and parameters*

Synchrotron-generated X-ray beams are tangentially emitted by relativistic electron bunches circulating in the storage ring of a synchrotron radiation facility. The experiments have been carried out at the ID17 beamline of the ESRF. The ID17 X-ray source is a wiggler (a magnetic structure of alternating poles positioned on a straight section of the storage ring, consisting of a sequence of 21 poles, with a period of 150 mm) producing a wide continuous spectrum of photons with energy ranging up to several hundreds of kilo-electronvolts (keV). The X-ray beam is quasi-parallel (horizontal divergence of 1 mrad, vertical divergence <0.02 mrad) with a quasi-laminar shape (maximum beam size at the sample position 40 mm horizontal x 1 mm vertical). After filtering with a sequence of beryllium (1.5 mm), carbon (1.5 mm), aluminum (1.5 mm) and copper (1.0 mm), the polychromatic radiation spectrum entering the treatment room had a mean energy of approximately 100 keV(Crosbie et al. 2015). The beam was spatially fractionated into an array of rectangular microbeams of variable width size by means of a multislit collimator(Bräuer-Krisch et al. 2009). The X-ray fluence produced by the wiggler determined an entrance dose rate of around 16,000 Gy/s, allowing the deposition in a fraction of second of doses of hundreds of Grays along the microscopic planes.

*Irradiation geometry*

An array of seven (7) parallel microbeams depositing an incident dose of 360 Gy was orthogonally directed over the bone vault onto the left somatosensory cortex (1 mm AP and 1 to 3.5 mm in ML direction). The microbeam size at the sample position was 103 μm and the center-to-center spacing was 412 μm at the collimator position providing a nominal mediolateral coverage of the irradiated cortex of 2.4 mm. The microbeam length in the anteroposterior (AP) direction was equal to 3 mm and the beam comb was positioned starting at 0.5 mm and ending 3.5 mm from the 0 bregmatic virtual horizontal line.

*Image-guided beam delivery*

Xylazine-Ketamine anaesthetized rats were placed vertically and fixed by ear bars and teeth on a home-made Plexiglas stereotactic frame and placed on a Kappa-type goniometer (Huber, Germany), by which the rat could be translated and rotated in front to the fixed horizontal X-ray microbeam array. To guide irradiations, low dose high resolution radiographs were acquired by vertically scanning the sample in front of the attenuated polychromatic X-ray beam (wiggler magnetic field set at 0.32 T) and by recording images using a 2048 x 2048 pixel FreLoN CCD camera (Coan et al. 2006), placed about 5 m downstream the sample. Radioscopies allowed for identifying the Bregma and therefore to center microbeams on the desired target.

For irradiations, the beam height was defined by a 520 μm wide tungsten slit, placed at 1 m upstream the animal. The dose was delivered by vertically moving the Kappa goniometer, at a speed inversely proportional to the dose to be delivered. Each irradiation lasted less than 1 s. The vertical irradiation field was determined by the opening-closing of fast shutters, located 7 m upstream the rats and synchronized with the movement of the Kappa goniometer. All movements were remotely controlled and irradiation values were preset by the operator before the treatment. The animal immobility (granted by the deep anesthetic state) during exposure was checked by 3 high resolution cameras located in the control hutch. The incoming spatially non-fractionated dose was measured using an ionization chamber and the mid-valley doses were calculated by Monte Carlo simulations (Siegbahn et al. 2005).

*Dose profiles*

The dose profile into the target consists of an alternation of peaks and valleys, i.e., of high doses along the microbeam path and low doses in the spaces between them. The reference valley dose is normally assumed as the minimum dose in between two microbeams. The valley dose must not exceed the maximum dose tolerated by the normal tissues traversed by the radiation. Its value depends on several parameters including the photon energy, the number of microbeams, the microbeam width and height, the center-to-center distance, as well as the depth, shape, and composition of tissues proximal to, within, and distal to the target(Siegbahn et al. 2005). The dose deposited by the microbeams (360 Gy) refers to the peak entrance dose measured at 3 mm in depth; it is constantly decreasing in the target with a half value layer of ~4.5 cm in water. The Monte Carlo calculated valley doses were as low as 4.4 Gy at 3 mm depth with a kind of build-up reaching 5.3 Gy at 1 mm in depth. A logarithmic scale the two different profiles for the peak and the valley dose in the target is shown in Fig. 1E-F.

**Behavioral tests**

The behavioral tests consisted of three stages. Freely walking in a transparent plastic arena (45x65 $cm^2$, having 10 cm high vertical borders), observations and estimates of the walking patterns, their speed and correct paw placement on the floor of the arena floor (Movies 1-4 exemplary for the four experimental groups), time estimates of paw retraction-latencies during von Frey tests and hot-plate tests of the left posterior limb in all the four experimental groups, i.e. CR, CCI, CCI-MB and CR-MB rats. All tests were performed one or two weeks after the MB treatment.

*Walking Patterns*

The walking schemes have been evaluated just before the preparatory surgery and on the second and the final day before electrophysiology. Besides the walking scheme irregularities and the sensory threshold anomalies, no other evident anomalous behavioral sign has been observed both in feeding or sleep rhythms but for the natural weight loss in the first days after the generation of the CCI model (Millan et al. 1987).

*Sensory tests: thresholds and latencies*

The released stimuli were applied by von-Frey and Hot plate tests (Gårdmark, Höglund, and Hammarlund-Udenaes 1998) measuring the paw retraction latencies to the application of light threshold and noxious stimuli. Specific signs and symptoms such as spontaneous postural guarding and paw withdraw during gait combined with allodynia and hyperalgesia in sensory tests were all observed and measured. Before proceeding with the recording sessions, a final behavioral validation test evaluated that all the signs of chronic pain were clearly detectable. The motor patterns were checked by a walking motor scheme test, showing the postural rhythms and patterns of paw placing during walking. In addition, mechanical allodynia has been measured by stimulating with von Frey filaments (Laird and Bennett 1993) the plantar aspect of the treated paw (well accessible after placing the rat over a metal grid). Specifically, animals were housed in a special cage with a grid floor that permits the assessment of sensory thresholds by applying successively increasingly rigid plastic filaments each bending at specific increasing forces when applied on the rat hind limb palm.

Measures of paw withdraw latencies were then collected for all the rats every other day with double or triple tests in short time (5 minutes) intervals. Heat hyperalgesia was evaluated by the Hot plate test, again estimating by an automatic system the paw withdrawal latencies on the heated plate at 51°C (Gårdmark, Höglund, and Hammarlund-Udenaes 1998). This test was repeated only on the second day after the preparation surgery and on the last day before the electrophysiological experiments, a strategy to avoid heat induced damages of the paw surface. The walking schemes have been evaluated just before the preparatory surgery and on the second and the final day before electrophysiology. Besides the walking scheme irregularities and the sensory threshold anomalies, no other evident anomalous behavioral sign has been observed both in feeding or sleep rhythms.

**Electrophysiological experiments: Neuronal and EEG recordings**

Acute electrophysiology recordings of the somatosensory thalamocortical complex were performed in all the four groups of experimental animals (CR, CCI, CCI-MB, CR-MB) by means of couples of microelectrode matrices about one or two weeks after the MB treatment.

*Microelectrode Electrophysiology Surgical Preparation*

Twenty-eight male albino rats (Sprague-Dawley, Charles River, Calco, LC, Italy, 270–350 g) were randomly chosen out of a larger set of the animals. The rats underwent preliminary barbiturate anesthesia (50 mg/kg ip) for the surgical experimental preparation. The trachea was cannulated to gain a connection to the anesthesia-ventilation device. A 16-gauge butterfly was then positioned in the root of the lateral tail vein to grant intra-surgical or intra-experimental access to drug intravenous delivery. A surgical opening was then done over the head skin, with the removal of the skin, the *galea capitis* muscles and of their fibrous insertions over the parietal bones. The periosteum was then delicately scraped off from the parietal, frontal and occipital bones after lidocaine infiltration. The rats were then placed in a stereotaxic frame and the animal were paralyzed by intravenous Gallamine thriethiodide (20 mg/kg/h) injection and connected to the respiratory device delivering (1stroke/s) an Isoflurane® 2.5% 0.4 to 0.8 l/min in Oxygen 0.15–0.2 l/min gaseous mixture. Curarization was maintained stable throughout the whole experiment by Gallamine refracted injections.

*Electroencephalogram Surgical Field*

During the experiment the anesthesia level was continuously monitored by 4 EEG recording leads and 1 reference lead, placed, respectively in a fronto-occipital row (in stereotaxic coordinates(Paxinos and Watson 1998) referenced to the bregmatic point, where *F*, *P* and *O* refer to Frontal, Parietal and Occipital where *F* + 1.2 mm AP-2.2 mm LL; *P1*: − 1.2 mm AP, 2.5 LL; *P2*: -3 mm AP, 2.7 mm LL; *P3*: -5.5 mm AP, 2.7 mm LL) contralateral to neuronal recordings. The reference lead, was placed on the occipital bone posterior to the Lambda reference point (*O*: − 2 from Lambda, 2 mm LL). In order to ease the placement of the EEG leads, five partial bone holes have been trephined at those reference points and five external leads have been then placed in the bone cradles. These were connected to the recording system by a home-made plastic pre-built frame with 5 golden pitch spring-test probes with concave heads vertically inserted in the plastic frame in stereotactic array. A hypertonic saline cream was used to ameliorate the electric interface between the leads and the bone. During the experiment, the EEG recordings continuously monitored the anesthesia level. For the analyses, we selected preferentially the EEG data from the second derivation placed over the sensory cortex mirroring the contralateral somatosensory primary cortex where the neuronal recordings were obtained (A.G. Zippo et al. 2016).

*Neuronal Electrophysiological Recordings*

For the electrophysiological neuronal recordings, two holes were drilled on the skull of 3 $mm^2$ for the cortical and the thalamic matrix accesses. The holes were drilled centered respectively on the cortical access centered around a reference point at − 1.5 mm AP and 2.5 mm ML. The thalamic access hole was centered at − 6 mm AP and − 2.5 mm ML. We simultaneously recorded spiking and local field potential activities by two microelectrode matrices from the thalamic ventro-postero-lateral complex nuclei (VPL) and the primary somatosensory (S1) cortex. The neuronal electrophysiological recordings have been, obviously, obtained from the cortex contralateral to the surgical lesioned and stimulated paw; the two microelectrode matrices of extracellular Pt-Ir electrodes were framed in 3 × 3 arrays of single shanks, inter-tip distance 150–200 mm, tip impedance 0.5–1 MΩ (FHC Inc., ME, USA). The coordinates have been estimated from a Stereotaxic Atlas (Paxinos and Watson 1998). In detail, the two matrices were placed at − 1.2 mm AP, 2.6 mm LL and − 6 mm AP, 2.5 mm LL for the somatosensory cortex and the thalamic nuclei respectively. The cortical matrix was inserted 400 μm deep at the superior border of the fourth granular layer and then slowly advanced, by an electronically controlled microstepper (PI Instruments, Germany), at 10 μm steps. This strategy was used for probing the responses of local neurons to exploring sensory light cotton tip light stimuli on the contralateral posterior paw, until clear responses were evident and repetitive on at least six out of the 8 recording microelectrodes of the matrix within the final depth of 600 μm. The thalamic regions were targeted with a postero-anterior slant of 25° of the matrix to avoid spatial interferences with the cortical matrix. This obliged to recalculate the depth by a simple geometric correction to the usual measure. The probe was advanced electronically by a second electronically driven microstepper (PI Electronic, Germany) from a starting depth of 4700 μm down to 5800 μm (50 μm steps, quickly driven to avoid excessive tissue damage).
Fast thalamic and cortical responses to light tactile stimuli with a brush-test on the sciatic innervation field (the plantar aspect of the left hind limb) were the anatomo-functional acceptance criteria for starting signal acquisition. All the experimental blocks were organized with periods of ongoing activity recordings lasting around 20 min. and not more, to preserve at most the data homoscedasticity in the additional stable conditions of gaseous anesthesia.
After a cycle of spontaneous and stimulated activities was completed, we repeated twice the original recording series. Then we advanced in depth the electrodes, 20 μm and 50 to100 μm respectively for the cortical and the thalamic matrix ensembles, to reach an adjacent recording region, then checking again with the test stimulus the responsiveness of the newly recorded regions. In positive cases, we repeated the recording cycle as above. We recorded from five to six stations in progressive steps for each animal.
For signal amplification and data recordings a Cheetah Data Acquisition Hardware was used (Neuralynx, MT, USA, sampling frequency 32 kHz). Electrophysiological signals were digitized and recorded with bandpass at 6 kHz and 0.1 Hz for spikes and Local Field Potentials and at 475 Hz–1 Hz for the EEG. The data stored were analyzed off-line both by Matlab and by locally developed software.
Two experimental conditions were considered: the resting state and the sensory stimulus state in which we applied a validated technique (Antonio G. Zippo et al. 2013; A.G. Zippo et al. 2014) to generate evoked spatially confined and fine-tuned tactile stimulations of the rat hindlimb. Briefly, smoothed wooden tips mounted vertically on the belly of woofer dustcaps were fast displaced (1-2 mm runs) under electronically regulated pulses. The tip touched a surface of some 100 $\mu m^2$ on mapped points on the volar surface of the treated paw, namely the fingertip pads and the thenar and hypothenar pads successively in sequences of stimulus trains. The trains were released with random intervals (minimum interval 1 s in order to avoid learning or habituation phenomena (Antonio G. Zippo et al. 2013; A.G. Zippo et al. 2014)).

### Microscopic Analyses

Histological examination of the regions of interest of the Somatosensory Cortex allowed for identification of the MB traces within the cerebral matter. Nissl staining and Osmium or Carbon coated Scanning Electron Microscopy (SEM) imaging have been made. Immuno-histochemical analyses have been done in alternate slices to conventional and SEM imaging to analyze local damages by immunofluorescence technique.

*SEM analyses*

Textural and morphological observations (secondary electron images) were performed by means of a Tescan FE-SEM (Mira 3XMU-series), equipped with an EDAX energy-dispersive spectrometer. The operating conditions were: 15 kV accelerating voltage, around 40 mA beam current, different working distance (reported in each photo), counts of 100 s per analysis. The measurements were processed using the EDAX Genesis software and semi-quantitative data obtained using the ZAF correction (where Z is the atomic number correction related to stopping power of the element, A is the absorption correction, F is the fluorescence correction). SEM imaging has been used at diverse magnifications from 2x to 300x to analyze the MB-generated tissue scars both in the skull bone and within the brain tissue. The bone tiles analyzed at SEM were the samples extracted during the electrophysiological experiments after trephination of the skull vault to clear the cortical area for the recordings. The brain histological slices were prepared along common protocols with rats under deep barbiturate anesthesia perfused with heparinized saline followed by 10% formalin solution perfusion. A primary gross section of the removed brains was made isolating a brain block (by a flat blade) with two coronal cuts, at bregmatic landmarks 0 and at -3.5mm, along the antero-posterior axis. The histological slicing was then performed on these blocks after inclusion in paraffin. The 5-6 and 10 µm thick brain slices have been obtained from each region of interest of all the brains. Histological slices prepared in alternate fashion (one slice for common histology, the next for SEM analyses) were placed on separate slides and treated either with histological recipe from Nissl staining or with osmium or carbon. In this last condition, a carbon source (a pure graphite rod) was mounted in a vacuum system between two high-current electrical terminals. When the carbon source was heated to its evaporation temperature, a fine stream of carbon was deposited onto the histologic sample. Osmium preparation was made with traditional methods by using a 4% aqueous solution and histological samples placed for 12 h in the solutions and then left to dry under chemical hood. Traces of brain MB irradiation were then looked after at low magnification survey of the brain slices with special focus on the cortical left side until the recognition of the thin scars aligned in medio-lateral arrangements. Then, progressive magnification within the cut trajectories was performed, analyzing both the morphology and the chemical element composition of the beam involved tissue as well as of untouched tissue regions (SEM HV 7.0 kV; Det. In Beam).

*Bone and Brain MB signs in microscopy*

Scans have been done on the skull bone involved in the MB irradiation to understand if bone lesions were detectable after the irradiation. Clear tiny traces of the MB irradiation were well evident throughout the whole cortical thickness (Magnification 71x, Field of View 4.07 mm 15 keV, Dt: SE). The comparison with the accompanying image in conventional Nissl staining of the same region of the same cortical sample, in a slice 20 um posterior to that obtained in SEM imaging along the fronto-occipital axis, confirms the coherence of the MB in their length extension.
Quantitative estimates of elements C, N, O, Na, Mg, Al, Si, Os, S, Ca were detected in all the scans in coherent amount of the brain tissue.

**Preliminary data analyses**

The data stored were analyzed off-line both using Matlab and by locally developed software. The neural firing rates had a mean of 31.4 Hz with standard deviation of 26.8 Hz. After the recordings, the LFPs were downsampled to 0.5 kHz. After filtering and downsampling, the spike contamination of LFP signals was null avoiding further spike removal techniques. The spikes were extracted and sorted by using the *Wave_clus* MATLAB toolbox. Sorted cells with average rates below 4 Hz and above 100 Hz were excluded from the analysis. Furthermore, neurons resulted from sorting which had improbable inter-spike-interval distributions were discarded as well. Recorded neurons were uniformly distributed over the recording matrices and every electrode showed distinct neural activity. At the end of this process, we collected a total of 3584 neurons, 128 ± 17 in each experiment, out from the set of the acquired signals.
The timestamps of spike occurrences were represented by binary sequences where 1's labeled a spike. We considered time bins of 1 ms, thus avoiding occurrence of multiple spikes within the same bin.
Finally, we split each sequence into fixed-length (250 ms) overlapping windows thus obtaining an ordered set of equal length windows. In order to discriminate among groups of recording by the firing rates, we verified the intraclass consistency of the recorded spiking activity within each experimental condition. To this purpose, we performed a one-way analysis of variance on the four sets of experiments. No significant difference within each class was observed.

**Functional connections by spike-train similarities**

Interactions between neurons can generate very complex, time-delayed and asymmetric patterns especially in the thalamocortical circuitry. In this work, we proposed a framework successfully applied in a similar context wherein spike trains were modeled by Markov stochastic models (Antonio G. Zippo et al. 2013).
We used the function Normalized Compression Similarity (NCS), formally defined as: given that $x$ and $y$ are two spike trains, the NCS is equal to

$$NCS(x,y) = \frac{C(x \cdot y) - min\big(C(x), C(y)\big)}{min\big(C(x), C(y)\big)}$$

where the C function represents the compressed sequence length and · is the sequence concatenation operator (e.g. 0101·101 = 0101101). If $NCS(x,y)$ is close to 1, the sequences $x$ and $y$ are considered similar. If close to 0, the sequences are strongly dissimilar. We evaluated the NCS function on time windows (250 ms length) of the recorded spiking activity, assuming that relative high values of similarity corresponded to actual functional connections.

**Functional connections by LFP phase synchrony**

LFPs are low frequency signals reflecting a wide range of synaptic events. In this work, we investigated the synchrony of LFP phases originated in different recording sites during spontaneous and tactile evoked activities. We measured phase synchronies between two recorded LFP sequences (x and y) by the following function:

$$\gamma(x,y) = \left|\left\langle e^{i\left(arg\left(H(x)\right)-arg\left(H(y)\right)\right)}\right\rangle\right|$$

where $e$ is Napier's constant, H is the Hilbert Transform, *arg* is the argument function and $i$ is the imaginary unit. The Hilbert transform and the argument were computed with, respectively, the *hilbert* and the *angle* Matlab functions. When $\gamma(x,y)$ is equal to 1 (0), then x and y are perfectly synchronous (asynchronous).

**Complex Brain Networks**

By using the $NCS$ and $\gamma$ functions, we estimated the functional connections of the recorded neuronal networks. We first split each recorded sequence into 250 ms time windows and then we computed the adjacency matrix for all neurons or electrodes. The resulting matrices exhibited values in the unitary interval. The functional connections extracted from extracellular recordings are the counterpart of non-oriented graphs. We noticed that variable thresholds equal to a percentile of the weight distribution varying between 0.2 and 0.8 did not affect results and thus we selected the 75$^{th}$ percentile.

For the analysis of these graphs, we introduced a set of common statistics from the Complex Network Theory able to detect possible matches between the extracted graphs and prominent brain topologies (Table 1) (A.G. Zippo and Castiglioni 2016; Antonio G. Zippo et al. 2013).

From a functional perspective, brain networks can express two important information-processing features: information integration and segregation. Functional segregation recruits specialized processing within densely interconnected nodes (cliques). Functional integration combines information processed in distributed nodes or cliques. These network properties can be measured by two statistics: the clustering coefficient (C) and the characteristic path length (L). The former measures how close the neighbors of a node are to being a clique. The latter estimates the average shortest path length in the graph, i.e. how much the nodes are accessible. Both measures, implemented in a Matlab toolbox, were used for our network analyses (*clustering_coef_bu.m*, *charpath.m*).

Eventually, we analyzed networks that evolved in time, dropping and recruiting nodes, connections and networks from different experimental conditions. Such a methodology requires the discussion of potential issues. First, unconnected nodes were rare but could occur after adjacency matrices were binarized. For this reason, we removed graphs in which less than 99% of nodes were connected. Second, network statistics were applied on networks with different sizes (for spiking activity) because the recording sessions returned a variable number of active neurons. However, by analyzing the observed variance of network size we concluded that C and L couldn't be significantly affected by our network size changes. Significant changes appeared for synthetic networks that increased their size by orders of magnitude. However, we discarded graphs that were outliers (beyond 5$^{th}$ and 95$^{th}$ percentile) of the node, edge and density number distributions in order to obtain a better homogeneity (Antonio G. Zippo et al. 2013). In the work, we refer to these two conditions as *admissibility criteria*.

**Software Accessibility**

The computer simulator for the cortical network was derived from the "Vertex Simulator" (www.vertexsimulator.org) and the code is available at https://sites.google.com/site/antoniogiulianozippo/codes.

Bennett, G J, and Y K Xie. 1988. "A Peripheral Mononeuropathy in Rat That Produces Disorders of Pain Sensation like Those Seen in Man." *Pain* 33 (1). NETHERLANDS: 87–107.

Bräuer-Krisch, E., H. Requardt, T. Brochard, G. Berruyer, M. Renier, J. A. Laissue, and A. Bravin. 2009. "New Technology Enables High Precision Multislit Collimators for Microbeam Radiation Therapy." *Review of Scientific Instruments* 80 (7): 074301. https://doi.org/10.1063/1.3170035.

Coan, Paola, Angela Peterzol, Stefan Fiedler, Cyril Ponchut, Jean Claude Labiche, and Alberto Bravin. 2006.

“Evaluation of Imaging Performance of a Taper Optics CCD `FReLoN’ Camera Designed for Medical Imaging.” *Journal of Synchrotron Radiation* 13 (3): 260–70. https://doi.org/10.1107/S0909049506008983.
Crosbie, Jeffrey C., Pauline Fournier, Stefan Bartzsch, Mattia Donzelli, Iwan Cornelius, Andrew W. Stevenson, Herwig Requardt, and Elke Bräuer-Krisch. 2015. “Energy Spectra Considerations for Synchrotron Radiotherapy Trials on the ID17 Bio-Medical Beamline at the European Synchrotron Radiation Facility.” *Journal of Synchrotron Radiation* 22 (4): 1035–41. https://doi.org/10.1107/S1600577515008115.
Gårdmark, M, A U Höglund, and M Hammarlund-Udenaes. 1998. “Aspects on Tail-Flick, Hot-Plate and Electrical Stimulation Tests for Morphine Antinociception.” *Pharmacology & Toxicology* 83 (6): 252–58. http://www.ncbi.nlm.nih.gov/pubmed/9868743.
Laird, J M, and G J Bennett. 1993. “An Electrophysiological Study of Dorsal Horn Neurons in the Spinal Cord of Rats with an Experimental Peripheral Neuropathy.” *Journal of Neurophysiology* 69 (6). UNITED STATES: 2072–85.
Millan, M J, A Członkowski, C W Pilcher, O F Almeida, M H Millan, F C Colpaert, and A Herz. 1987. “A Model of Chronic Pain in the Rat: Functional Correlates of Alterations in the Activity of Opioid Systems.” *The Journal of Neuroscience : The Official Journal of the Society for Neuroscience* 7 (1): 77–87. http://www.ncbi.nlm.nih.gov/pubmed/3027278.
Narula, Vaibhav, Antonio Giuliano Zippo, Alessandro Muscoloni, Gabriele Eliseo M. Biella, and Carlo Vittorio Cannistraci. 2017. “Can Local-Community-Paradigm and Epitopological Learning Enhance Our Understanding of How Local Brain Connectivity Is Able to Process, Learn and Memorize Chronic Pain?” *Applied Network Science* 2 (1). Springer International Publishing: 28. https://doi.org/10.1007/s41109-017-0048-x.
Paxinos, G, and C Watson. 1998. “The Rat Brain Atlas in Stereotaxic Coordinates.” *San Diego: Academic*.
Siegbahn, E.A., E. Bräuer-Krisch, J. Stepanek, H. Blattmann, J.A. Laissue, and A. Bravin. 2005. “Dosimetric Studies of Microbeam Radiation Therapy (MRT) with Monte Carlo Simulations.” *Nuclear Instruments and Methods in Physics Research Section A: Accelerators, Spectrometers, Detectors and Associated Equipment* 548 (1): 54–58. https://doi.org/10.1016/j.nima.2005.03.065.
Zippo, A.G., and I. Castiglioni. 2016. “Integration of $^{18}$FDG-PET Metabolic and Functional Connectomes in the Early Diagnosis and Prognosis of the Alzheimer’s Disease.” *Current Alzheimer Research* 13 (5).
Zippo, A.G., S. Nencini, G.C. Caramenti, M. Valente, R. Storchi, and G.E.M. Biella. 2014. “A Simple Stimulatory Device for Evoking Point-like Tactile Stimuli: A Searchlight for LFP to Spike Transitions.” *Journal of Visualized Experiments*, no. 85. https://doi.org/10.3791/50941.
Zippo, A.G., S. Rinaldi, G. Pellegata, G.C. Caramenti, M. Valente, V. Fontani, and G.E.M. Biella. 2015. “Electrophysiological Effects of Non-Invasive Radio Electric Asymmetric Conveyor (REAC) on Thalamocortical Neural Activities and Perturbed Experimental Conditions.” *Scientific Reports* 5. https://doi.org/10.1038/srep18200.
Zippo, A.G., M. Valente, G.C. Caramenti, and G.E.M. Biella. 2016. “The Thalamo-Cortical Complex Network Correlates of Chronic Pain.” *Scientific Reports* 6. https://doi.org/10.1038/srep34763.
Zippo, Antonio G., Riccardo Storchi, Sara Nencini, Gian Carlo Caramenti, Maurizio Valente, and Gabriele Eliseo M. Biella. 2013. “Neuronal Functional Connection Graphs among Multiple Areas of the Rat Somatosensory System during Spontaneous and Evoked Activities.” Edited by Olaf Sporns. *PLoS Computational Biology* 9 (6). United States: Public Library of Science: e1003104. https://doi.org/10.1371/journal.pcbi.1003104.
